# Using a Homeogram to Detect Sleep in Free-living Animals

**DOI:** 10.1101/2021.10.14.464397

**Authors:** Matt Gaidica, Emily Studd, Andrea E Wishart, William Gonzalez, Jeffrey E Lane, Andrew G McAdam, Stan Boutin, Ben Dantzer

## Abstract

1. Sleep is appreciated as a behavior critical to homeostasis, performance, and fitness. Yet, most of what we know about sleep comes from humans or controlled laboratory experiments. Assessing sleep in wild animals is challenging, as it is often hidden from view, and electrophysiological recordings that define sleep states are difficult to obtain. Accelerometers have offered great insight regarding gross movement, although ambiguous quiescent states like sleep have been largely ignored, limiting our understanding of this ubiquitous behavior.
2. We developed a broadly applicable sleep detection method called a *homeogram* that can be applied to accelerometer data collected from wild animals. We applied our methodology to detect sleep in free-ranging North American red squirrels (*Tamiasciurus hudsonicus*) in a region that experiences drastic seasonal shifts in light, temperature, and behavioral demands.
3. Our method characterized sleep in a manner consistent with limited existing studies and expanded those observations to provide evidence that red squirrels apply unique sleep strategies to cope with changing environments.
4. Applying our analytical strategy to accelerometer data from other species may open new possibilities to investigate sleep patterns for researchers studying wild animals.

## Introduction

Sleep is a ubiquitous animal behavior, such that—with rare exception—*you know it when you see it.* Sleep is typified by a common body position, reduced responsiveness and mobility, rapid reversibility, and a homeostatic rebound following deprivation (Ungurean et al., 2020). Sleep also serves common functions; from reptiles to birds to mammals, it usually places an animal in a temporary environment that reduces the risk of predation, decreases thermoregulatory and metabolic demands, and optimizes access to shared resources (Anafi et al., 2019; Eban-Rothschild et al., 2017). It is therefore unsurprising that studying sleep in unnatural or augmented environments, such as the laboratory, eliminates many of the challenges and pressures that influence sleep patterns on daily to evolutionary timescales (Rattenborg et al., 2017).

Tools and efforts to quantify sleep in the wild are few compared to those for the laboratory (S. S. Campbell & Tobler, 1984; Capellini, Barton, et al., 2008; Capellini, Nunn, et al., 2008; Gravett et al., 2017; Lesku et al., 2006, 2009; Massot et al., 2019; Rattenborg et al., 2017; Stuber et al., 2015). Also, definitions of sleep are based upon neural-electrophysiology data, making it an inherently invasive endeavour (Aulsebrook et al., 2016; Stickgold & Walker, 2009). However, multi-axis accelerometers that quantify movement may have some utility in this realm (Walch et al., 2019). After all, sleep and active behavior are usually antithetical (Ruckebusch, 1972).

Efforts to characterize sleep using body motion have, arguably, been more vigorously pursued towards applications related to human health, notably in the past decade due to wearable consumer and medical electronics. This comprises an immense literature concerning accelerometer-based sleep methods—some proprietary—that are tailored towards stereotyped, human sleep phenotypes (Altini & Kinnunen, 2021; Ancoli-Israel et al., 2003; Cole et al., 1992; Enomoto et al., 2009; Nakazaki et al., 2014; Paquet et al., 2007; Sadeh, 2011; Sadeh et al., 1994; Sazonov et al., 2004; van Hees et al., 2018). However, underlying themes emerge in data analysis techniques with regard to sleep. Ground-truthed data either from field observations or laboratory animals can ‘label’ accelerometer data (i.e. behaviors visually identified in-sync with sensor data) which strengthens the ability of algorithms—such as supervised machine learning— to build models that directly map actigraphy features to behavior (H. A. Campbell et al., 2013; Hammond et al., 2016; Tatler et al., 2018). Auxiliary sensors can also be used as features as well (e.g., collar temperature), but these are often not standard in type, sampling frequency, precision, or deployment method across disciplines. The resulting sleep algorithm may take the form of a ‘black box’ (Shuert et al., 2018), decision tree (Khanh et al., 2016), or resolve optimal parameters to pre-defined equations (Chimienti et al., 2016; Patterson et al., 2019; R. P. Wilson et al., 2006), which can be used with new, unlabelled data.

Despite relative success in characterizing sleep with labelled data, much of the data coming from wild animals in quiescent states is unlabelled due to the difficulty of observing animals in their natural sleep habitat (e.g., underground, in nests, or otherwise hidden in many terrestrial animals) (Bedford et al., 2021; Brown et al., 2013). Furthermore, many species have ill-defined sleep patterns, such that, despite a rich suite of seemingly similar tools for studying sleep using accelerometry, few can be applied to wild animals since they rely on a priori knowledge of sleep patterns or impose arbitrary, species-specific rules to their classification procedure (Chimienti et al., 2016).

Here, we develop a robust, tunable, and physiologically consistent method of characterizing sleep-wake patterns using accelerometry alone that we call a *homeogram*. Our approach uses an alternative method of reconciling unlabeled behavioral data by viewing it in terms of the underlying biological processes that it represents. It is inspired by the two-process model of sleep proposed by Borbély (1982) where sleep pressure is formed by the interaction of a homeostatic process and circadian process. Our homeogram takes a Borbélyian approach to modeling the overarching circadian rhythm based on a few rules and incorporates ubiquitous features of sleep, such as the fact that it has both intensity and inertia, all while facilitating the possibility of short sleep-wake episodes that occur day or night (Figure 1). The result is a sleep estimation (as a probability) that can be binarized into asleep and awake states with minimal code and language-independent mathematical transformations.

**Figure 1.**
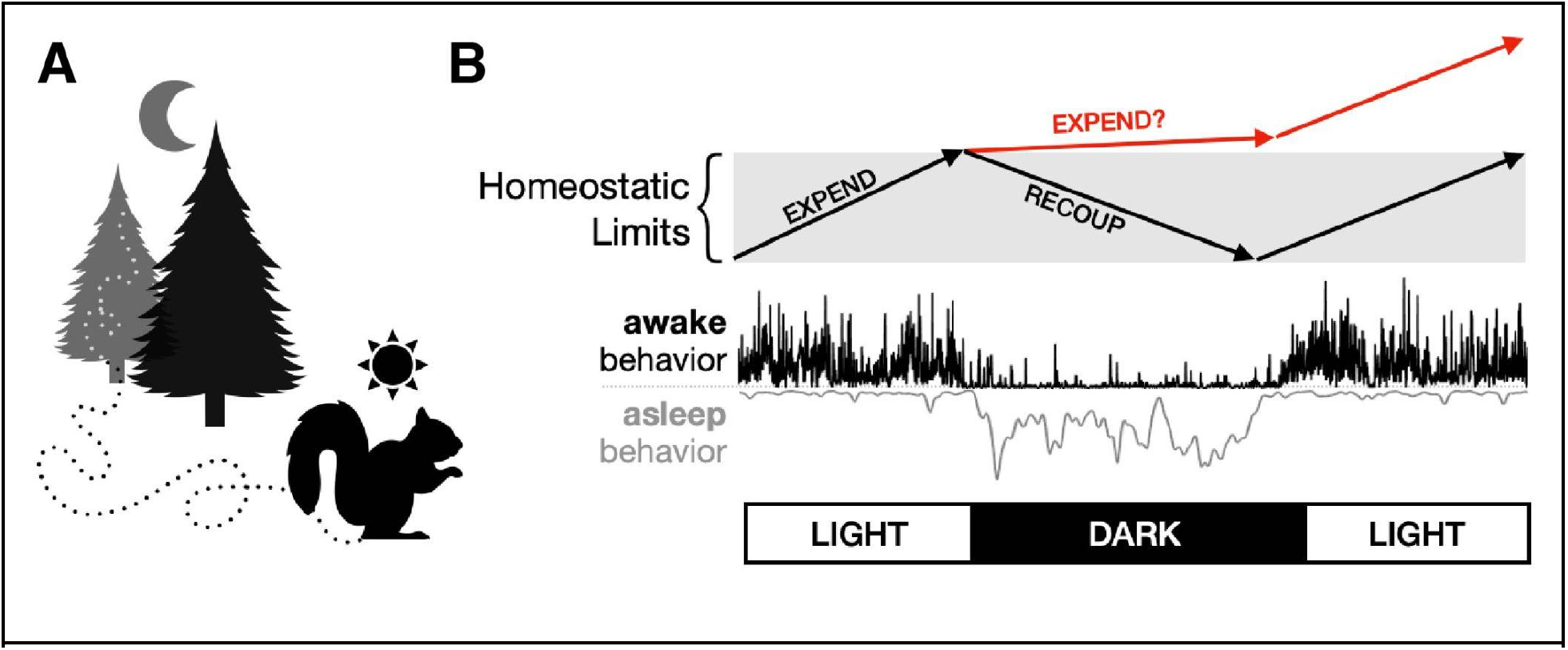
The Missing Link in Accelerometer Data. (A) Accelerometers typically focus on quantifying active behaviors, but not quiescent ones like sleep. (B, top) Because they quantify movement, an accelerometer is inherently ‘positive-only’ data. From a physiological perspective, this reflects expenditure but not recuperation, and therefore needs to be modified if those data need to be interpreted in a homeostatic model (i.e. where expenditure and recuperation are roughly equalized over a period of time). (B, bottom) These mock data incorporate a conceptualization of homeostatic processes actually occurring in an animal; an ideal method to detect sleep should recognize that sleep is a behavior with amplitude and inertia, rather than simply the absence of movement.

To test the effectiveness of our homeogram method to characterize sleep, we compiled accelerometer data from five years of study of North American red squirrels (*Tamiasciurus hudsonicus*) in Yukon, Canada. Red squirrels in this region are subject to high northern latitudes, which have a daylight difference of over 13 hours between summer and winter, representing a unique methodological and physiological challenge. Like many animals, despite a great deal of understanding regarding red squirrel active behavior (Dantzer et al., 2013, 2016; Fisher et al., 2019; Klugh, 1927; Studd et al., 2016, 2020), little is known about sleep patterns in general or how they use sleep to cope with changing seasonal demands.

Below, we first describe how to construct a homeogram from raw acceleration data and validate the algorithm using nest attendance (based on collar temperature). Next, we use the binary form of the homeogram to generate sleep statistics and provide a gestalt view of how red squirrels sleep from daily to yearly time scales. Finally, we investigate the rhythmicity of red squirrel sleep and whether or not perturbations in ambient food availability affect sleep patterns. Altogether, we use a homeogram along with conventional analyses of sleep to show how researchers can engage sleep research in free-living animals without new or invasive data collection (i.e., using accelerometer data).

## Application

### 1. Homeogram Construction

The homeogram takes its name from the *actigram*, which is a time-series histogram indicating activity magnitude as recorded by an accelerometer or other activity sensors (e.g., running wheel encoder, infrared beams, piezoelectric plates). In contrast, a homeogram attempts to use similar data to estimate expenditure (as positive numbers) and recuperation (as negative numbers). Thus, a homeogram is designed to reflect the fact that sleep has both a circadian process and homeostatic value. That is, it is assumed an organism maintains an energetic equilibrium over time (Refinetti, 2007). Under that premise, the basic chain of processing a signal from raw accelerometer data to a mutually exclusive asleep and awake classification is straightforward (Figure 2A). This begins by computing overall acceleration (OA), which is the sum of the absolute value of all accelerometer axes after subtracting a running median to remove static acceleration (Equation 1); for our data, we used a median window of 91 seconds, which was empirically determined based on a sensitivity analysis developed by Shepard (2008) and which we previously applied to red squirrels (Studd et al., 2019). This has similarities to the original development of overall dynamic body acceleration (ODBA, R. P. Wilson et al., 2006) but does not exclude the possibility of low accelerometer sampling rates (≤ 1 Hz) that do not accurately delineate static versus dynamic movements. In fact, we further downsampled our data by a factor of 60 using a moving mean over the original 1 Hz data to an effective rate of once-per-minute (0.0167 Hz) to support data exploration and hypothesis testing without the need of an advanced or complex computing workflow.

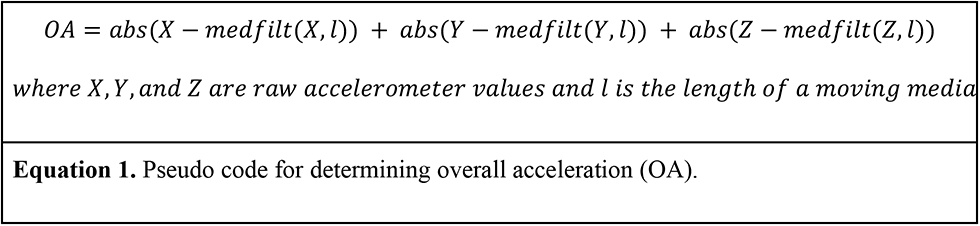

**Figure 2.**
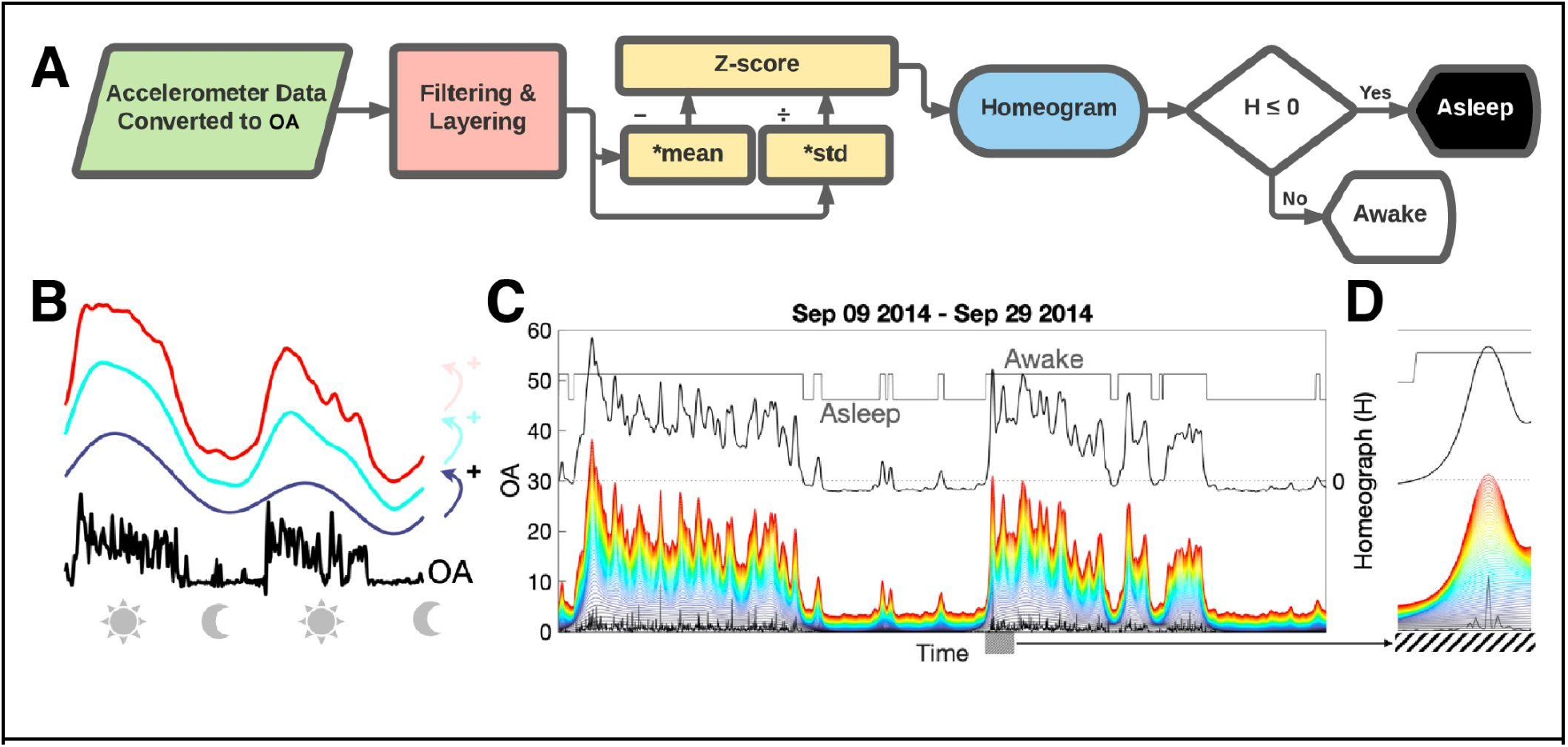
Constructing the Homeogram and Detecting Sleep. (A) Accelerometer data is first converted to overall acceleration (OA; see Equation 1) to remove the dependency on accelerometer orientation. Filters are applied to OA and layered (i.e. summed) progressively then the signal is Z-scored using the mean and standard deviation of quiescent periods (*mean and *std in figure) to standardize the signal across individuals. The result is the time varying Homeogram (H), where H ≤ 0 is asleep and H > 0 is awake. (B) The layering process from OA of a diurnal animal (black, bottom) showing the progressive filters that are layered on top of each other; blue is the first filter layer, cyan is the sum of the first and second, and so on until all layers are summed (depicted in red). (C) Actual resolution of filtering and layering from a free-living squirrel collected September 9-24, 2014. The top-most line (red) is what enters the Z-score process, resulting in the Homeogram (black line, right y-axis). The binary asleep-awake classification is shown atop. (D) A zoomed-in view of C for the hatched area.

The homeogram uses progressively summed layers of a locally regressed quadratic filter (LOESS) which has superior qualities to a moving mean or Gaussian window because of its local sensitivity to changes in OA amplitude (Figure 2B). The first filter length is the size of day length or night length, whichever is least. Therefore, the ‘base’ of the homeogram has similar smoothing qualities for long and short days and conceptually represents the ebb and flow of expenditure and recuperation—or sleep-wake activity—of a diurnal or nocturnal animal. Subsequent layers are summed using progressively smaller windows at 1-hour intervals (24 filters total) which add detail to the circadian base. Cruder determinations of sleep-wake cycles may choose to use fewer filter layers, whereas polyphasic phenotypes from animals that sleep in many short bouts throughout the day may benefit from eliminating the base layers, and hibernators from adding them (revisited in the Discussion).

The final step to create the homeogram is to normalize it, which is useful to ensure all data—which may be coming from different sources—has similar characteristics before the asleep-awake classes are applied. Theoretically, this makes the specific recording properties of the accelerometer hardware (or activity sensor) unimportant. We accomplished this by first estimating where quiescence exists after the filtering and layering stage by applying a Z-score transformation (Figure 2C, D). The mean and standard deviation of segments with a Z-score less than zero were used again on the original signal to create the homeogram. Note that this process results in a time-series where the mean is not zero, and therefore, the homeogram is not a Z-score as it is conventionally defined. Conceptually, this approach treats quiescent data segments as periods that estimate accelerometer ‘noise’ and which should be relatively consistent across all data. Although, we acknowledge that interventions that could enhance animal movement during these times (e.g., disturbances, increasing rapid eye movement sleep) might need more careful handling if they are important characteristics of the final analysis. Once the homeogram is created, it can be used as a time-varying probability that the animal is asleep (where negative numbers indicate a higher probability), or binarized about zero where positive values are awake, and values less than or equal to zero are asleep. The binary form can be easily used to identify sleep transitions (awake → asleep → awake), where the gradient of the signal is non-zero.

### 2. Homeogram Validation: Nest Attendance

External temperature sensors affixed to an animal (such as on a collar or harness) can be a useful surrogate for determining if an animal is inside of a nest or burrow given that the temperature should rise for endothermic animals (Studd et al., 2016) and act as a gatekeeper for classifying active, waking behavior (Studd et al., 2019). While it might seem like a useful feature to also gate sleep classification, some animals—such as red squirrels—are often observed outside of their nest sleeping or basking in the sun, rendering the collar temperature data a problematic determinant of sleep itself, nonetheless, a good estimate of overall nest use patterns (Steele & Koprowski, 2003). Collar temperature also approximates qualities of thermal-based validation measured previously used to assist in delineating sleep from wake (Loftus et al., 2021). As a first step in understanding if a homeogram concurs with these expectations, we quantified the relationship between nest attendance and asleep behavior from a multi-year dataset of red squirrels (see Figure 3) by extracting the binary determination of each, where nest attendance was based on collar temperature and asleep-awake state based on our homeogram approach using accelerometry.

**Figure 3.**
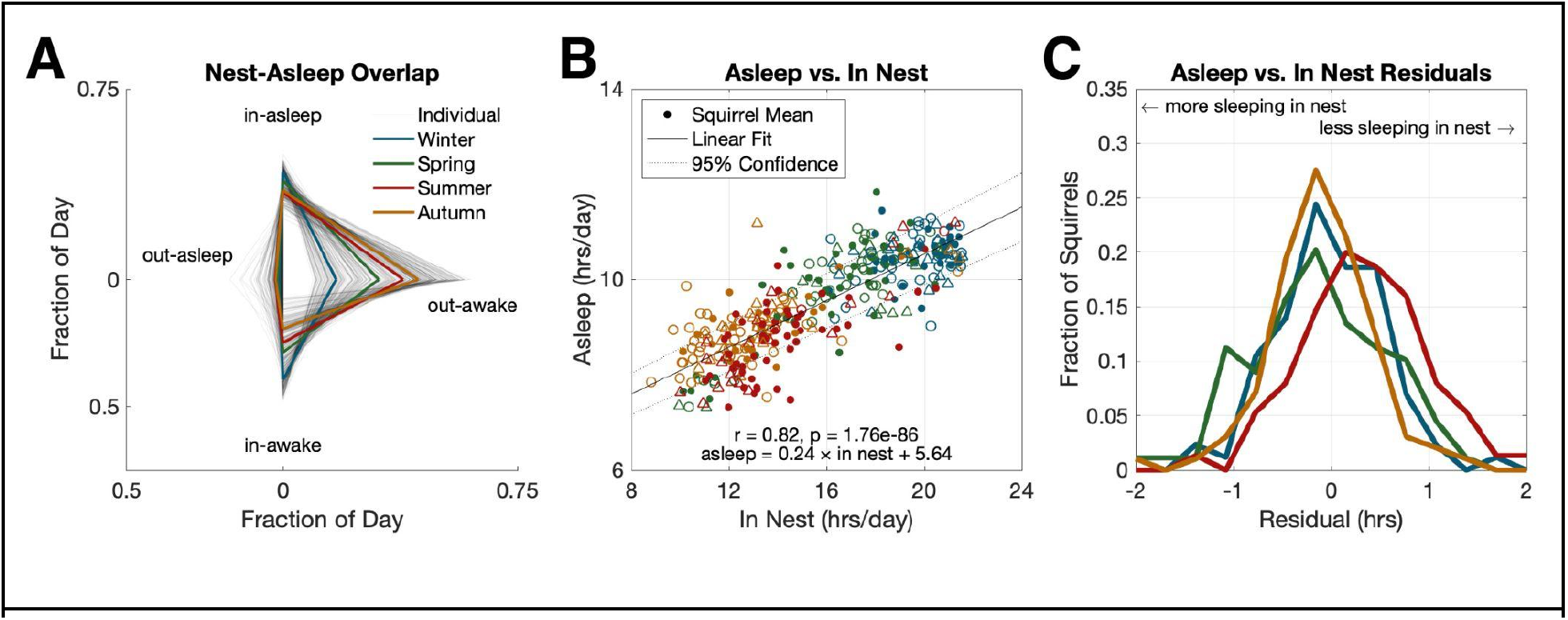
Sleep-wake states and nest attendance. (A) Fraction of day (i.e., time) spent between sleep and nest states (345 recording sessions representing 218 squirrels) over four seasons (mean values in colored lines which apply to all subplots). In-nest and out-nest were determined using collar temperature data (Studd et al., 2019). Season was determined by the mean recording day (see breakdown below, Table 1). (B) The relationship between time spent Asleep (determined using Homeogram, originating from OA) and In Nest (determined by collar temperature). Squirrels with 1 recording session are solid dot markers (●), 2 recording sessions are open triangle markers (△), and 3 or more recording sessions are open circles (○) colored by season. Pearson linear correlation values are shown along with the best-fit line equation. (C) Residuals from B for each season highlighting seasonal differences in times spent in nest sleeping.

**Table 1.** Sleep statistics for North American red squirrels in Yukon, Canada derived from homeogram using accelerometry data. Sleep transitions characterize awake to asleep or vice versa.

|  | Winter | Spring | Summer | Autumn | All |
| --- | --- | --- | --- | --- | --- |
| Day of Year | 311-35 | 36-127 | 128-218 | 219-309 | 1-366 |
| Recording Sessions | 84 | 88 | 75 | 98 | 345 |
| Unique Squirrels | 71 | 67 | 66 | 68 | 218 |
| Day Length (hrs.) | 6.55 ± 0.79 | 12.44 ± 2.52 | 18.21 ± 0.81 | 12.37 ± 2.47 | 12.38 ± 4.52 |
| Sleep per day (hrs.) | 10.46 ± 1.90 | 9.91 ± 2.16 | 8.73 ± 2.01 | 8.73 ± 1.70 | 9.51 ± 2.10 |
| Sleep in daylight (hrs.) | 0.73 ± 0.87 | 1.88 ± 1.67 | 4.29 ± 2.04 | 0.88 ± 1.45 | 1.96 ± 2.14 |
| Sleep in darkness (hrs.) | $9.73 \pm 1.82$ | $8.03 \pm 1.75$ | $4.44 \pm 1.07$ | $7.85 \pm 1.24$ | $7.55 \pm 2.56$ |
| Total sleep transitions | $28 \pm 7$ | $24 \pm 8$ | $19 \pm 7$ | $14 \pm 6$ | $22 \pm 9$ |
| Sleep transitions in daylight | $3 \pm 3$ | $6 \pm 4$ | $13 \pm 6$ | $4 \pm 4$ | $7 \pm 6$ |
| Sleep transitions in darkness | $25 \pm 6$ | $18 \pm 7$ | $6 \pm 4$ | $11 \pm 5$ | $16 \pm 9$ |

We found the expected pattern of nest-asleep behavior (Figure 3A). Most notably, Winters are characterized by an increased amount of time in the nest—both awake and asleep— whereas animals transition to being more outside-and-awake towards Autumn (Studd et al., 2020). The presence of some sleep outside of the nest is consistent with the observation that sleep may not always occur in the nest. Furthermore, nest attendance does not directly equate to sleep but is highly correlated (Figure 3B). Plotting the residuals from this relationship shows a left-shift for Autumn (Figure 3C), suggesting a more efficient use of the nest for sleep (i.e., squirrels sleep a higher proportion of time when in the nest). Summer exhibits a mirrored pattern, suggesting less efficient use of the nest for sleep, whereas Winter and Spring are neutral in this regard (i.e. normally distributed).

To further investigate these features and control for repeated observations of the same squirrel, we used a linear mixed effects model that included sex and season as fixed effects and squirrel identity as a random effect (Table S2-4). Squirrels spent significantly more total time sleeping (in nest and out of nest) in the Winter and Spring months compared to Summer and Autumn (Tukey’s p <0.0001 for all pairwise comparisons), whereas there was no difference in total time spent sleeping between Summer and Autumn (Tukey’s p = 0.57). Overall, there was no significant effect of sex on total time sleeping (p = 0.136) or sleep outside of the nest (p = 0.151), but females slept less in the nest (p = 0.0259).

In summary, the homeogram meets our expectations both visually and analytically as a method to extract asleep-awake behavior using accelerometry. Even movements in the limited volume of a nest are reliably identified as wakefulness which is an important distinction from analyses that treat nest attendance as sleep or rely on ‘first nest exit’ and ‘last nest entry’ as a proxy for sleep patterns (see Wassmer & Refinetti, 2016, Fig. 7). While we benefited from prior work that established an algorithmic nest classification from collar temperature (in essence, labeling our data), future implementations of the homeogram do not rely on these multisensor data, but may benefit from them. Furthermore, when we tested the homeogram against a widely used accelerometer algorithm (Sadeh et al., 1994) on human data, we found 95.54% agreement (Figure S2), confirming that it is generally accurate, but benefits from the features described here.

### 3. Putting Sleep Data in Context

Once sleep is binarized using the homeogram, compiling a set of sleep statistics is trivial (Table 1). Briefly, we split our data from red squirrels into four equally sized seasons based on mean day length (i.e., time between sunrise and sunset) between Spring and Autumn; this resulted in two seasons where behavior could be directly compared since light availability is nearly equal. Sunrise, sunset, and solar noon were derived using a sunrise equation (US Department of Commerce, 2021) that uses the latitude and longitude of our study location of red squirrels in Yukon, Canada (60° 45’ 16.0344”, -137° 30’ 42.4074”) and daylight savings time (DST) was corrected by applying the local time zone in MATLAB. Note, universal standard time (UTC) is advised when DST might not be coordinated between sensors and observers since it is not affected by DST.

We compiled mean asleep values around a 60-day window for each day of the year (Figure 4). This visualization can also be conceptualized as the sleep probability of a red squirrel at any day and time. These results demonstrate the robust plasticity between sleep phenotypes based on season for red squirrels. Winter is highlighted by relatively inconsistent sleep, which accounts for a lower level of sleep in darkness, as well as a small peak during the day where naps (sleep in daylight)—or at least, deep rest—may be taking place. Despite the importance of Spring as it relates to reproduction, it sticks out as a period of a gradual transition to the long Summer days, where pre- and post-noon naps sprout, consistent with our finding that a large amount of sleep occurs during the day in Summer (see Table 1). Autumn is more well-defined and sharply delineated than Spring, likely a result of the stricter demands on daily activity, and perhaps, nightly recovery.

**Figure 4.**
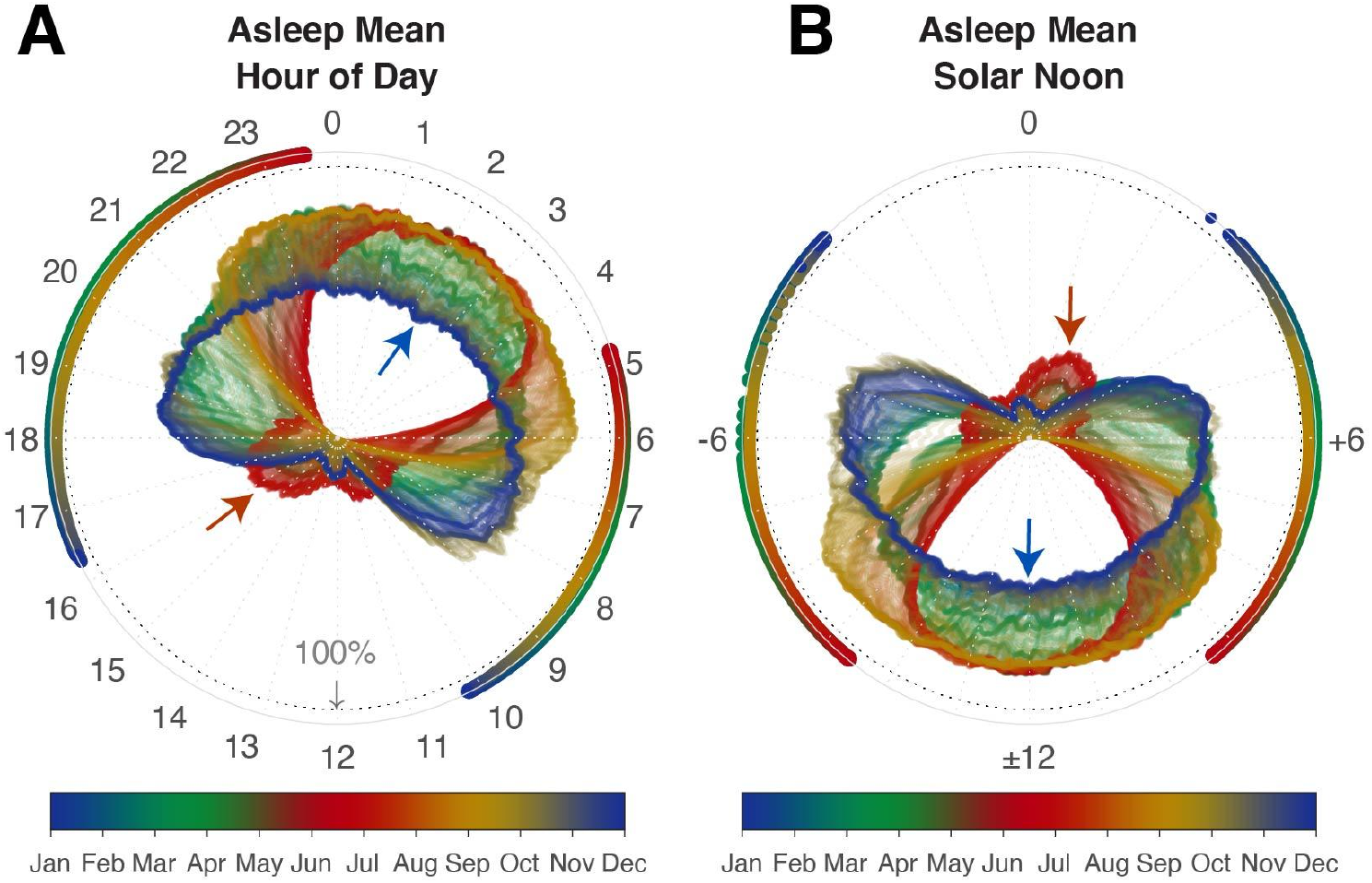
Fraction of red squirrels asleep. Daily red squirrel (345 recording sessions, 218 squirrels) sleep patterns were determined for each day of the year (denoted by color). A 60-day, centered mean is presented around the 24-hour solar cycle represented as the hour of day (A) and relative to solar noon (B). For both plots, the light dashed line labeled ‘100%’ represents where all squirrels would be asleep; the center, inner radius is where no squirrels are asleep. Outer colored marks denote sunrise and sunset. In both plots, the red arrow highlights putative naps emerging in the daytime, and the blue arrow shows where sleep is less consistent across recordings (i.e., ∼50% of squirrels are sleeping at this time).

Figure 4 can be easily converted to a time-series plot to understand the nature of sleep phenotypes for each season (Figure 5). These results show how season determines a sleep phenotype, rather than day length, by comparing Spring and Autumn. The strong transitions at sunrise and sunset of Autumn and the small variance in daytime sleep suggest a ‘seize the day’ approach to this important season where squirrels spend their time caching cones from white spruce trees (*Picea glauca*) to serve as a food source for the coming winter (Fletcher et al., 2010; McAdam et al., 2019).

**Figure 5.**
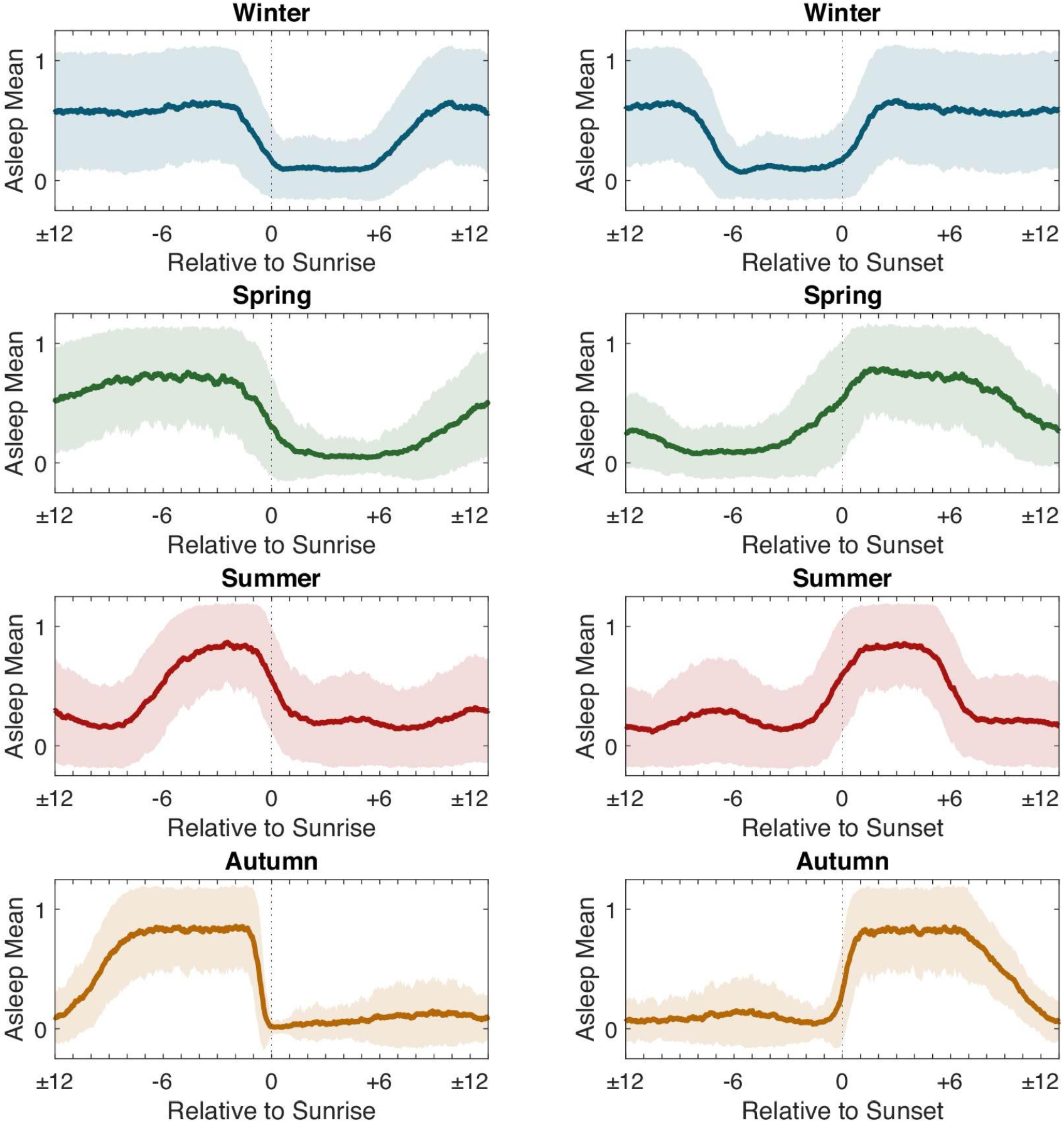
Asleep Mean and Standard Deviation. Binary asleep-awake states were determined from the homeogram (where asleep = 0, awake = 1) and used to determine the time-varying mean and standard deviation relative to sunrise (left column) and relative to sunset (right column).

### Rhythmicity Analysis

Rhythmicity is a measure of how consistent a behavior (e.g., sleep) pattern is over time. Rhythmicity can be a useful metric to compare across seasonal or interventional conditions since entrainment is a fundamental component of organismal biology and can correlate with overall health and condition (Mistlberger & Rusak, 2011). In brief, the rhythmicity of a signal can be quantified using an autocorrelation analysis. This method scrubs a signal across itself and calculates a correlation between those lagged signals. Thus, at a lag of 0, the autocorrelation is always 1 and symmetrically extends to each side for positive and negative lags up to the length of the data. Therefore, rhythmic data will exhibit positive peaks at locations *n*-cycles from a lag of 0. To showcase its utility, we hypothesized that a rhythmicity analysis would further support Autumn as the *bourgeonner* of highly regulated sleep-wake behavior in red squirrels, as well as resolve the ultradian rhythms (i.e., naps) embedded in Winter and Summer (see Figure 4).

We modified the binary awake-asleep classification from the homeogram approach described above such that sleep was negatively signed (−1) and awake was positively signed (+1) so they would have the properties of a standard unit oscillation. Each recording session was then autocorrelated and averaged for each season (Figure 6, left). As we expected, primary rhythms were present at 24-hour intervals across all seasons. We calculated the rhythmicity index (RI) as the third peak (including the 0-lag condition) (Dowse, 2009; Levine et al., 2002) which revealed Autumn as the most rhythmic season for awake-sleep patterns. The adjacent season of Winter has an RI of nearly half, altogether consistent with our prior results suggesting that sleep is most variable in Winter.

**Figure 6.**
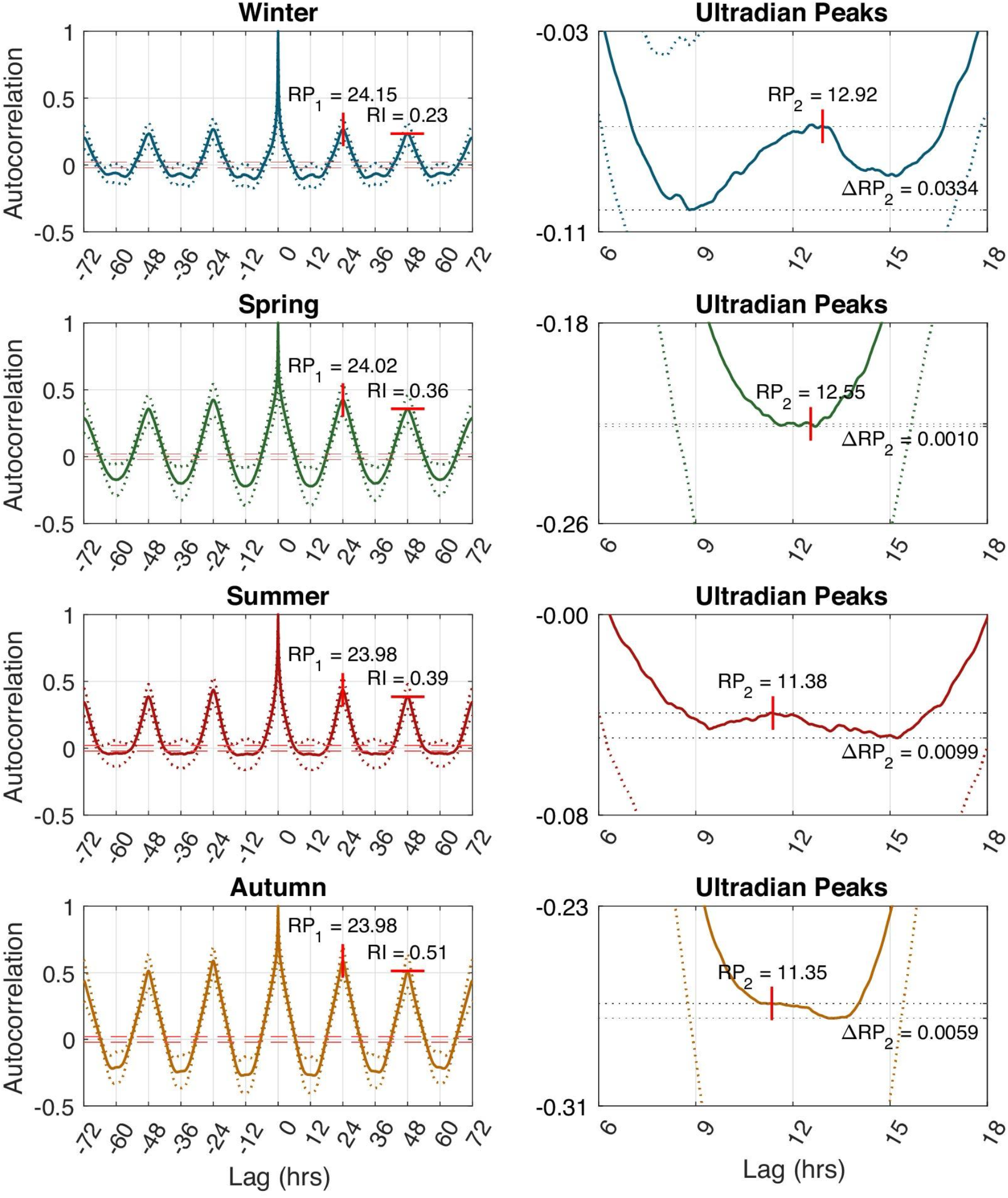
Sleep Rhythmicity. The autocorrelation of sleep quantifies the period and intensity of its underlying rhythm. (Left column) The primary rhythm period (RP_1_) is the circadian rhythm (i.e., roughly 24-hours). The Rhythmicity Index (RI) quantifies the rhythm intensity as the third peak. Values between the red dashed lines (95% confidence) are considered ambiguous. (Right column) The secondary rhythm period (RP_2_) is the ultradian rhythm. ΔRP_2_ is the greatest local change in rhythm intensity. For both plots, standard deviations are dashed lines.

**Figure 7.**
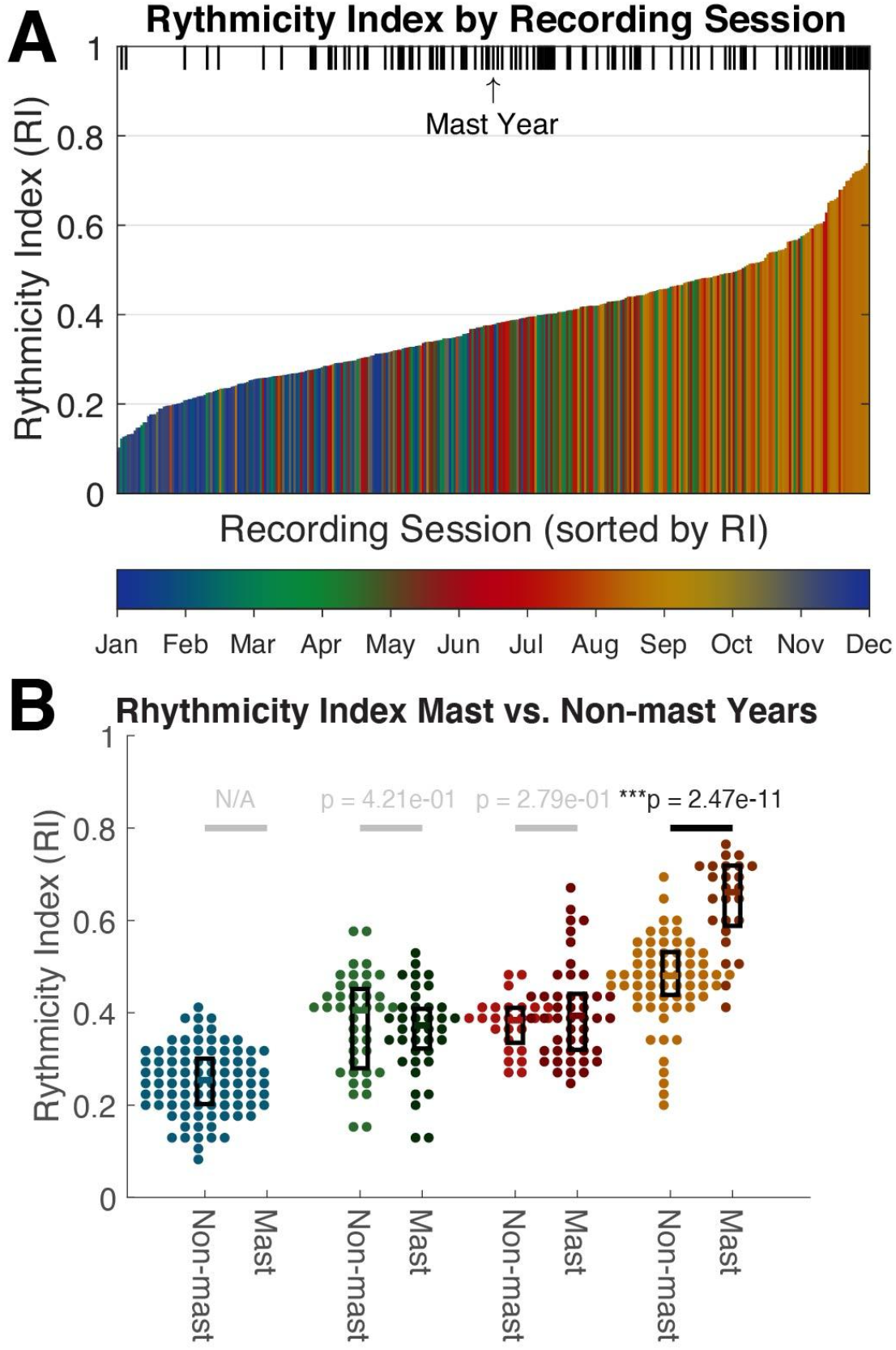
Rhythmicity Index by Recording Session and Season. (A) Recording sessions from each season are color coded and sorted by rhythmicity index (RI, y-axis). Mast years, where food is super-abundant and physical demands are increased due to hoarding and caching are marked with a vertical pipe along the top. (B) Beeswarm plot of RI for each recording session (●) colored by season (from left to right: Winter, Spring, Summer Autumn) and organized by Mast and Non-mast years. P-values of the one-way ANOVA testing the null hypothesis that the means of the groups are equal is shown above each Mast vs. Non-mast season group. We did not have Winter mast data.

We then aimed to quantify potential ultradian (i.e., less than 24-hour) peaks by focusing on the autocorrelation centered about 12-hours (Figure 6, right). A purely monophasic circadian signal with sleep opposed to wakefulness would exhibit a strong negative correlation at 12-hours lag, whereas, if harmonics (i.e., ultradian rhythms) existed, the negative peak would be interrupted with positive correlations. We found ultradian peaks that were invariably unimodal, so we identified the single largest peak and quantified its magnitude over the local minima. Spring and Autumn exhibited minuscule ultradian rhythms, whereas Winter and Summer were the largest. For comparison, Winter’s ultradian rhythm amplitude was 22.3 times larger than Spring’s, 6.0 times stronger than Autumn’s, and 1.5 times stronger than Summer’s. The Winter and Summer rhythms were only slightly phase-shifted from the primary rhythm. Based on our previous data showing the potential for napping in Winter and Summer, we interpret this result as suggesting that when naps do occur, they are nearly perfectly out of phase with the primary sleep rhythm. For instance, red squirrels are most likely to be asleep around 3–4 A.M. and most likely to nap—180° out of phase—around 3–4 P.M.

### Awake-Sleep Patterns during Dynamic Environmental Conditions

A rhythmicity analysis based on the homeogram may be useful in discerning subtle sleep dynamics across fluctuating environmental conditions. For example, under non-stressful situations the hypothalamic-pituitary-adrenal axis operates under circadian control, however, exposure to external stressors may result in central nervous system activation that disrupts a healthy sleep schedule (Vargas et al., 2018). We were able to test this by investigating the change in RI during years in which white spruce trees produced a superabundance of cones available to squirrels in late Summer and early Autumn, which are known as mast years (Kelly, 1994). At our study site in Yukon, red squirrels primarily consume the seeds contained in white spruce cones, which they collect and cache underground in the Autumn of each year (Fletcher et al., 2010, 2013). Mast years occur every 4-6 years during which squirrels exhibit an extended breeding season, increased reproductive output, and substantial increases in cone hoarding behavior (Boutin et al., 2006; Dantzer et al., 2020; McAdam et al., 2019). Our hypothesis was that this naturally occurring condition would preferentially increase RI for Autumn of mast years, when red squirrels are busy hoarding and caching cones, since a stricter sleep presumably reduces the transition cost (i.e., latency to and from waking behavior) and increases overall productivity.

We compiled the RI for all recording sessions and examined its relationship to non-mast (2015, 2016, 2017) and mast (2014, 2019) years, and season (Figure 7). Consistent with the autocorrelation data, Autumn accounted for the largest RIs, which also appeared to be associated with a mast year. When we compared the distribution of RIs for each season, we found that the only significant difference was between Autumn non-mast and mast years (p = 2.95^-14^, one-way ANOVA), supporting our hypothesis that the increased demands of a spruce mast year enhance the rhythmicity of sleep-wake patterns.

### Sleep Transitions

It has been debated whether or not observed behavioral rhythms are the result of a ‘masking effect’ whereby discrete events, like sunrise and sunset, gate behavior rather than behavior being regulated endogenously (Rietveld et al., 1993). We hypothesized that sleep patterns were masked by sunrise and sunset, since daylight has a strong influence on the ability of a diurnal animal to forage and navigate, as well as visually identify threats. We used homeogram data to compile transition times in a 60-day window for each day of the year (similar to Figure 4) and plotted them aligned to sunrise and sunset or centered about solar noon (Figure 8). Both transition types have a subtle gradient across seasons. For example, as Autumn progresses, transitions to awake precede sunrise and follow sunset to a greater degree. When aligned to solar noon, the gradation largely disappears, not supporting our hypothesis concerning masking. Rather, this analysis suggests that the circadian clock is entrained to solar noon, when the sun passes the local celestial meridian and exerts the greatest amount of solar energy on earth. In other words, each sleep season appears to be determined by a solar-centered sleep quota rather than sunrise and sunset. Autumn and Spring can be compared as equivalents and exhibit large differences in sleep transitions, which supports our previous findings that sleep-wake behavior is plastic and may be based on behavioral demands or is indeed shaped through the feedback of those behaviors on the squirrel’s physiology.

**Figure 8.**
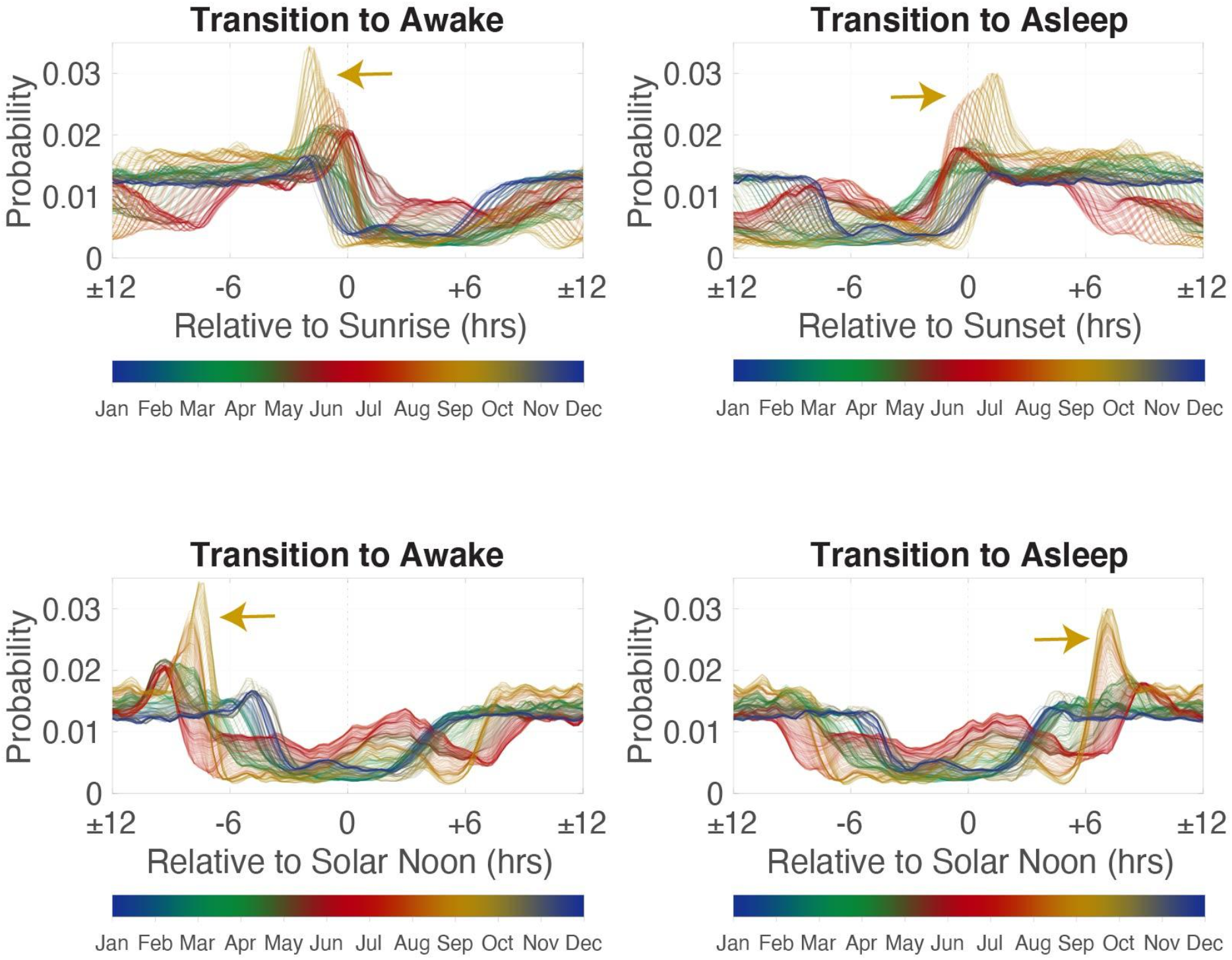
Sleep Transitions. Sleep transitions were calculated from the binary form of the homeogram and the probability for 60-day windows around a single day of year are plotted for transitions to the awake state (left column) and transitions to asleep (right column). The top row is transition probability relative to sunrise and the bottom row is transition probability relative to solar noon. The same sleep transitions (marked by arrows) are graded when relative to sunrise (top row) but become aligned relative to solar noon (bottom), namely in days of the Autumn season.

Sleep transition data extracted from the homeogram also enable sleep fragmentation and sleep bout analyses. That is, how often sleep is interrupted and the duration of each sleep bout, respectively. We hypothesized that sleep becomes less fragmented or ‘deeper’ in red squirrels in response to great physical/behavioral demands required to harvest and hoard spruce cones in the Autumn (Studd et al., 2020; see also Fletcher et al., 2010). We defined sleep duration as the period between a transition to asleep and transition to awake and confirmed that they can be described by a power law (Dimanico et al., 2021; Lo et al., 2002, 2004). Evidently, sleep duration distributions are significantly different between each season (p < 0.0001 for each pairwise comparison using Two-sample Kolmogorov-Smirnov test), where Winter sleep durations are shorter and Autumn sleep durations are longer (Figure 9). We next examined our data separated into mast (2014, 2019) and non-mast (2015, 2016, 2017) years to test if squirrels exhibited longer sleep durations in an environment of increased physical demands associated with caching cones (Boutin et al., 2006; Dantzer et al., 2020; Fletcher et al., 2010; McAdam et al., 2019; Studd et al., 2019). Despite not having mast year data for Winter, we found that Summer (p = 0.0288) and Autumn (p = 1.28^-20^) had significantly longer sleep durations in mast years than non-mast years. Furthermore, Summer mast sleep durations were less than Autumn non-mast sleep durations, and Autumn mast sleep durations were longer than all others (Figure 9 inset).

**Figure 9.**
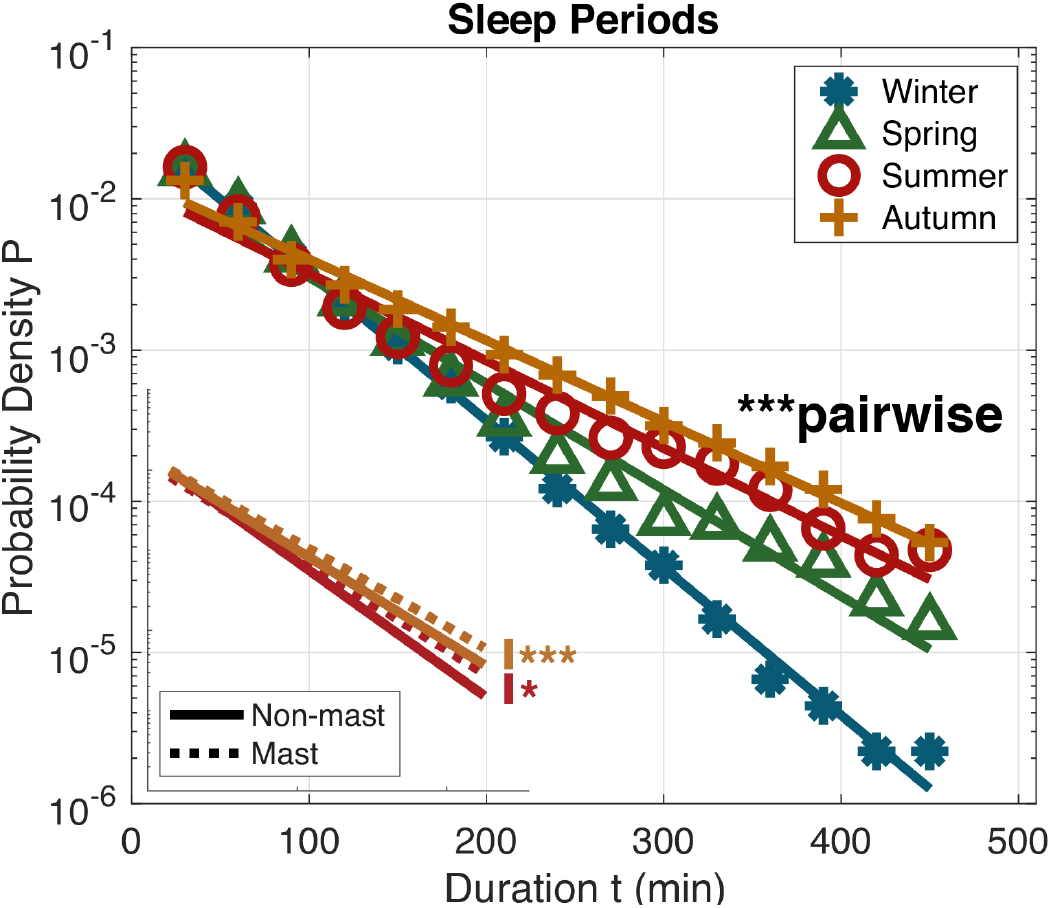
Seasonal Sleep Periods. Semilogarithmic plot for distributions of sleep durations. Raw binned (n = 20 bins) data are shown with markers with accompanying lines of best fit. All pairwise comparisons were significantly different. Inset shows Summer and Autumn Non-mast (solid) and Mast (dashed) best fit lines with the same axes as the main figure. Mast to non-mast comparisons were non-significant for Spring and mast data was unavailable for Winter. *p < 0.05, ***p<0.001 Two-sample Kolmogorov-Smirnov test.

Overall, sleep transitions are reduced in Autumn, which could simply be due to the increase in daily activity caused by caching cones. This is supported by the observation that sleep duration in darkness (but not daylight) is similar between Spring and Autumn (mast and non-mast; see Table 1). However, because of the Spring and Autumn day length equivalence, we could compare sleep transitions in the dark, which are far fewer in Autumn (Table 1). Most interesting is that the overall duration of sleep in Autumn is less in a mast year than in Autumn in non-mast years (Table S1). We again used a linear mixed-effects model to investigate the proportion of time spent sleeping in the nest (Table S5-6), which was not affected by sex (p = 0.1532). The lack of a significant interaction between mast year and season indicates that no matter mast status, the proportion of time spent sleeping in the nest in Autumn is significantly greater than all other seasons (p < 0.05) except Summer, where it is non-significant in non-mast years (p = 0.0755) but significant in mast years (p < 0.0001), likely because masting begins at the end of Summer as we defined it here.

In sum, the homeogram developed here allowed us to use accelerometer data to determine sleep rhythmicity, the prevalence of sleep transitions, and the duration of sleep periods. When applied to our data collected from red squirrels in Yukon, a unique model was formed by which squirrels sleep least in Autumn and even less in the Autumn of a mast year, but that sleep is more rhythmic and sounder (i.e., less fragmented and longer bouts), as well as more efficient from a nest-use perspective. Thus, we find support for the hypothesis that squirrels exhibit deeper (i.e., less fragmented) sleep in response to increased ecological and behavioral demands.

## Discussion

### The woods are lovely, dark and deep. But I have promises to keep, and miles to go before I sleep. ∼ Robert Frost

Motivated by the prevalence and utility of accelerometer deployment in ecological research and the relative paucity of insights into the growing field of sleep research, we developed the homeogram to translate accelerometer data into a sleep probability which is then easily binarized into asleep and awake classifications. Unlike many existing methods that require labelled data or complex rules to quantify sleep from accelerometer data, the homeogram is based on biological ‘first principles’, and therefore, we believe it is broadly applicable. Our method applied to data collected from free-living red squirrels shows how our method can be used to test hypotheses regarding when and how free-living animals sleep according to ecological challenges that they experience. We discovered that sunrise, sunset, solar noon, and behavioral demands interact with sleep demands. Furthermore, the unique features of Autumn in mast years where squirrels are collecting and caching food engender higher quality sleep based on its four pillars: depth, duration, continuity, and regularity (M. P. Walker, 2017). Below, we expand some of the major themes of our findings and methodological issues moving forward.

### Sleep and Homeostasis

Cannon defined the word *homeostasis* as “the tendency towards stability” in 1939, and nearly five decades later, Borbély integrated a homeostatic rhythm—interacting with a circadian rhythm—into his two-process model of sleep regulation (Achermann & Borbély, 2011; Deboer, 2018). In harmony, these two processes describe the coordination of endogenous and exogenous factors that should enhance the performance and fitness of individuals (Roth et al., 2010).

The hierarchy of mechanisms engendering homeostasis through sleep depends on its duration and intensity (Tobler, 2011) which can be defined multiple ways. From a behavioral perspective—the one that an accelerometer can see—sleep duration is time spent in the quiescent state, whereas intensity is the extent to which sleep is fragmented. Of course, hidden from the accelerometer are the neural electrophysiological states that are more nuanced. Firstly, sleep onset (and latency to this state after stillness) is typically demarcated by the first occurrence of slow-wave activity (SWA) indicative of non-rapid eye movement (NREM) sleep (Cano et al., 2008; Mckenna et al., 2008). This discrepancy often leads to accelerometers over-estimating sleep duration compared to electrophysiological measures (Ancoli-Israel et al., 2003). Therefore, the behaviorist might rely on a measure of sleep efficiency instead, derived by comparing the time spent still to the time spent in a nest, bed, or resting posture (Lutz et al., 2018). Secondly, the REM state offers the opposite problem in some mammals, especially infants, where the replay of active behaviors (which is normally gated from muscles) is temporarily allowed to pass through the spinal cord, resulting in high amplitude movements like twitching, trembling, and fluttering of the body and limbs (Louie & Wilson, 2001). We investigated the possibility of identifying such behavior by reconstructing the 3D trajectory of brief movements that were identified as wakefulness, but without a library of video observations or coordinated biopotential recordings, the task complicates rather than simplifies the interpretation of large-scale data. No more than 10% of brief awakenings in our data were deemed REM candidates. Taken together, there are certainly Type I and Type II errors present (Dekking, 2005), which may counterbalance, but remain a species-specific confound to accelerometry alone. Winnebeck et al. (2018) implemented a novel method of quantifying locomotor inactivity during sleep by amplifying the accelerometer data during putative sleep, which may broadly apply to distinguishing sleep-wake states. We also generally recognize the possibility that a larger proportion of the day nesting (e.g., Winter) increases the sleep false positive rate due to the limited range of motion that an animal has in a nest.

We used sleep fragmentation as a proxy for sleep intensity, which is electrophysiologically indicated by the presence of SWA (Mongrain & Dumont, 2007; Tobler, 2011). It is not uncommon for healthy individuals to experience fragmented sleep. Typically, brief awakenings are not remembered by humans in the morning because sleep inertia is too strong during those times for it to be registered in memory (Kenna, 2017). Sleep typically fragments for mammals at the completion of a sleep cycle, which is the stereotypical progression from a vigilant state to REM or NREM sleep, or the dreaming and homeostatic (i.e., deep sleep) states, respectively. However, it should be noted that REM sleep carries a potentially significant homeostatic value (Ephron & Carrington, 1966; Naiman, 2017; Pandey & Kar, 2018; Siegel, 2011). Nonetheless, when sleep cycles—and sleep architecture, in general—are disrupted, as evident by fragmented sleep, homeostasis is compromised (Reid & Zee, 2011). And despite the disparity between sleep cycle duration across species, being about 10 minutes for a small rodent and 90 minutes for a human (Zepelin & Rechtschaffen, 1974), fragmentation has similar deleterious effects across species (Toth & Bhargava, 2013). Fragmented sleep can be the result of a disease state or disorder (Jones et al., 1987; Kimoff, 1996; Mellman et al., 1995; Tafti et al., 1992) and effectively reduces optimal daytime vigilance and neurocognitive function (Bonnet, 1987; Stepanski, 2002). Indeed, fragmented sleep, which leads to daytime sleepiness, signals that the function of sleep has been inadequately met and remains functionally separate from sleep deprivation itself (Ramesh et al., 2009). In our red squirrel data, we found that Autumn had the least number of sleep transitions and the longest sleep periods, suggesting less sleep fragmentation. We found that these characteristics were more pronounced in years in which there was a superabundance of food that squirrels needed to collect and cache (mast years), where sleep duration is decreased (Table S1), reflecting a potentially adaptive mechanism to compensate for increased energy expenditure associated with cone collection and caching (Fletcher et al., 2012). Alternatively, activity itself and caloric intake are potent modulators of circulating hormones and circadian rhythms (Dallman et al., 2004; Dolezal et al., 2017; Laposky et al., 2008; Youngstedt et al., 2019), which may lead to better Autumn sleep. Autumn is also marked by enhanced sleep rhythmicity—which, at least in humans—is a desirable health trait (Gibson et al., 2009; Sulli et al., 2018). In fact, loss of circadian tone is a strong predictor of impending death in some, if not most animals (Pittendrigh & Minis, 1972; Wax & Goodrick, 1978)

### Ultradian Sleep Rhythms

Any rhythm occurring more than once per day is considered ultradian. This includes cycles of sleep states that normally come and go overnight (i.e., sleep architecture) down to heart rhythms. We were interested in the ultradian sleep rhythms that emerged in patterns characteristic of naps, most notably in Summer and Winter for red squirrels. Unlike most polysomnography, accelerometers have the advantage of recording continuously (but see Debellemaniere et al., 2018; Strangman et al., 2018), revealing quiescent patterns that exist outside the typical circadian sleep window (Cornelis et al., 2019; Miller et al., 2020).

The parsimonious answer to *why we nap* is to fulfill the additional requirements of sleep demand or a sleep quota. This is consistent with our findings in red squirrels in the summer months as they appear to sleep almost as much during the day than at night (similar to ground squirrels J. M. Walker et al., 1977). We suspect that the strong gating effect of sunrise and sunset (Wassmer & Refinetti, 2016) compresses sleep duration in Summer such that a nap must emerge, just as a tire tube might bulge at its weakest point under immense pressure; therefore, *when we nap* might be the more interesting question. Despite Summer naps appearing in the afternoon in red squirrels, our rhythmicity analysis suggests that the predominant nap forms a circasemidian rhythm occurring roughly 180° out of phase (∼12 hours) relative to the major, or core sleep period (Broughton, 1998). The reason for this may have to do with the fact that SWA, or homeostatic sleep, is correlated with the previous time spent awake. Thus, it may be advantageous to partition sleep bouts perfectly out of phase to minimize the accumulation of homeostatic pressure. This strategy can be extended to polyphasic sleepers that capitalize on frequent sleep sessions to engage SWA on a nearly continuous basis. One theory posits that every time an organism falls asleep, some process related to recuperation might be activated, allowing sleep benefits to accrue independently of sleep duration (S. Wilson, 1995). It is also known that, for monophasic diurnal animals, the most SWA occurs in major sleep with minimum body temperature, but for minor sleep, it is with maximum body temperature, which is very likely to follow solar noon (S. Wilson, 1995). Therefore, major and minor sleep sessions may be spaced to ideally reduce homeostatic pressure in coordination with ambient and internal body temperatures (Lavie, 1986).

The other reason *why we nap* is to conserve energy and could account for red squirrel naps in the Winter. While increased sleep variability across individuals in Winter likely dilutes the effect in the aggregate, our rhythmicity analysis suggests Winter ultradian rhythms are the strongest of all seasons. Because North American red squirrels do not hibernate or engage in torpor (Brigham & Geiser, 2012), they have high thermoregulatory demands throughout the year, which may be met by increasing time spent in the nest (by staying warm) while benefiting from the reduced energetic demands bestowed by sleep (Cerri & Amici, 2021; Schmidt, 2014). The relatively active morning and evening schedules during Winter likely serve a mixture of nutritional and territorial demands for the red squirrel. This interpretation is not mutually exclusive with the possibility that biological cascades increase sleep demand in the Winter. For example, exposure to acute stressors can promote NREM sleep and also trigger non-shivering thermogenesis (Harding et al., 2020) which may have net positive effects for an animal in extremely cold conditions, especially if nutritional resources are scarce. The frequency of sleep transitions in red squirrels is also highest in Winter. While this may be explained by a higher false-positive rate of sleep detection when an animal spends more time confined to a nest, frequent awakenings may also safeguard red squirrels from entering REM sleep—which occurs at the tail end of mammalian sleep cycles—where the control of body temperature is poikilothermic (i.e. minimal) (Carskadon & Dement, 2011).

### Method and Study Limitations

We implemented a motion-based sleep state classification that was tuned to identify sleep, but likely made Type I and Type II errors, which is potentially unsolvable using accelerometer data alone. In general, there is a tendency to overpredict the sleep state and underpredict the waking state (Sazonov et al., 2004). The lack of video or electrophysiology makes it impossible to delineate quiet rest from sleep or sleep movements from wakefulness in our data. In laboratory nests, red squirrels shift positions often and exhibit REM-like body twitching (Gaidica and Dantzer, unpublished data), although it is unclear if those movements occur during electrophysiological sleep. We are currently investigating methods to address such questions (Gaidica & Dantzer, 2020). A common attempt to circumvent these errors is to impose additional sleep-wake rules onto the primary algorithm (Webster et al., 1982). For example, “after at least 4 minutes scored as wake, the next 1 minute scored as sleep is recorded wake,” is one of many rules that result in marginal gains in algorithmic performance (Cole et al., 1992). These rules, which have evolved into more complex methods over time, still suffer from being empirical or trained on specific species and conditions, narrowing their applicability.

We did not focus on the source of variation between and within individuals, although our data included both sexes over many years. Investigating sleep-wake patterns by sex (W. Zhang & Yartsev, 2019), age (Steinmeyer et al., 2010), life-history stage (Wassmer & Refinetti, 2016), the prevalence of predators (White & Geluso, 2007), or climatic variation (Helm et al., 2013) are likely fruitful future directions. In red squirrels, day length and mean ambient temperature are positively correlated with overall activity levels, however, we do not have granular data regarding the light at, or temperature inside a given nest (but see Studd et al., 2016). Therefore, we cannot exclude the possibility that microclimates (e.g., the rise and fall of ambient temperature, winter snowfall, cloud cover, nest orientation) could also serve as a strong gate of wakefulness, although, we suspect ambient temperature is a proxy for behavioral requirements of a particular season for red squirrels (Efrat et al., 2019; V. Y. Zhang et al., 2019). We do know that lactating red squirrels select nests with varying insulative qualities to both reduce heat loss and dissipate heat (Guillemette et al., 2009) and the potency of thermal cues should not be underestimated, as they appear to underlie hormonal changes and circadian physiology of heterotherms which determine when to enter and exit hibernation (Lyman, 2014; Williams et al., 2016). Indeed, hibernation itself is difficult to treat with our algorithm without modification. While accelerometers are a good tool to characterize the entrance, maintenance, and exit of hibernation, their data not only require inspection in their raw form but are troublesome to interpret, since hibernation is not sleep (Blanco et al., 2016). Species-specific phenology is likely a necessary investment for the researcher wishing to apply accelerometry to hibernating animals (Williams et al., 2016) as it can be an ambiguous state of adapted inactivity. In addition, augmenting accelerometers with light or thermal sensors is likely to improve interpretation efforts (Barnes, 1989; Krauchi & Deboer, 2010; Williams et al., 2011).

## Conclusion

Here we present the *homeogram* to determine sleep-wake patterns from unlabeled accelerometer data. Applying the homeogram to a red squirrel data set, we revealed a mosaic of sleep-wake patterns that are highly plastic and then investigated those patterns using analyses familiar to the sleep or circadian scientist. If applied broadly, the homeogram may help in elucidating the evolutionary history of sleep phenotypes. For example, the phylogenetic history of the red squirrel would predict that they are polyphasic sleepers (Campi & Krubitzer, 2010). However, as we demonstrate, the sleep-wake patterns—namely in Autumn—appear indifferent from that of humans and other monophasic sleepers, indicating that some animals may exploit a latent ability to engage in briefer sleep episodes (than say, humans) while remaining overtly diurnal based on season or behavioral demands.

Although sleep is often appreciated as an emergent quality of wakefulness, we should consider the possibility that *wakefulness is an emergent quality of sleep*. As such, sleep research requires new tools—such as the homeogram—to progress.

## Acknowledgments

We thank Agnes MacDonald for long-term access to her trapline, and to the Champagne and Aishihik First Nations for allowing us to conduct work on their traditional territory. This research was funded by the Translational Research Institute for Space Health through NASA Cooperative Agreement NNX16AO69A, grants from the Natural Sciences and Engineering Research Council of Canada (to S.B., J.E.L., A.G.M.), the Northern Scientific Training Program (to students of S.B., J.E.L., A.G.M.), the W. Garfield Weston Foundation, and the USA National Science Foundation (IOS-1749627 to B.D).

## Author’s Contributions

MG developed the methodology, performed the analyses, and wrote the supporting sections of the manuscript with assistance from BD. WG assisted with data processing. BD, EKS, and AEW performed data collection. All other authors provided resources to enable data collection and performance site availability/access as well as provided feedback.

## Data Availability

All data will be available on Figshare.

## Supplementary Material

### Data Overview

#### Data Collection

Between 2014 and 2020, we collected accelerometer data from 218 squirrels over 345 unique recording sessions as part of the Kluane Red Squirrel Project located in Yukon, Canada (60° 45’ 16.0344”, -137° 30’ 42.4074”). Accelerometers (models Axy2/Axy3/Axy4) were obtained from Technosmart Europe and weighed about 4 grams before packaging into a collar (∼1.7% of body mass: Studd et al., 2020). Red squirrels weighed roughly 250 grams. Collars were mounted ventrally (when only the accelerometer was fitted) or dorsally when a VHF radio transmitter was added (model PD-2C, Holohil Systems Limited, Carp, Ontario, Canada; 4 g). Squirrels were captured by live-trapping and briefly handled in a canvas bag to place accelerometer collars on the squirrels using established methods (Dantzer et al., 2020). Each squirrel is permanently and uniquely marked with metal ear tags, enabling individual identification, and the identity, sex, reproductive condition, and body mass of the squirrel was recorded each time it was captured. Further details on the study area and general data collection methods are provided in Dantzer et al. (2020) and collar design is provided in Studd et al. (2019).

#### Data inclusion

Every recording session was manually reviewed for recording quality and questionable data/devices were removed from the analyses. All females that were lactating, pre-pregnant, or pregnant were excluded. Any recording with less than a day of data (1440 minutes) was excluded.

### Mast vs. Non-Mast Sleep Statistics

### Linear Mixed Effects Regression Tables

### Comparing Homeogram to Conventional Sleep-wake Algorithm

We obtained 7 days of human axy data (with ambient light intensity) from Vincent van Hees (https://accelting.com/) and reshaped it to exactly match ODBA of our squirrel data. We applied the Sadeh algorithm (Sadeh et al., 1994) by modifying the empirically-determined first term, which draws a threshold for sleep-wake classification. In brief, the Sadeh algorithm is formed as, *P(sleep) = Threshold – Weighted parameters applied to actigraphy*, where *P(sleep) > 0 = sleep*. In Figure S2, run our homeogram algorithm alongside the Sadeh algorithm and find 95.54% agreement. Where the algorithms disagree appears isolated to spurious deviations in the accelerometer data. Since the homeogram has a circadian base filter, these deviations these deviations are weighted towards ‘asleep’ during the primary sleep period, and towards ‘awake’ during the primary wake period. In other words, the homeogram adds circadian gravity for spurious data that may not represent a behaviorally relevant shift in movement/behavior. Sadeh-like algorithms or simply increasing the smoothing factor lack this feature, as they are highly localized, whereas the homeogram has a global context. Importantly, the Sadeh algorithm requires 5 terms that must be determined a priori—making it potentially useful for controlled animals labelled data is available—whereas the homeogram only requires 1, which is optional.

---

**Figure S1.**
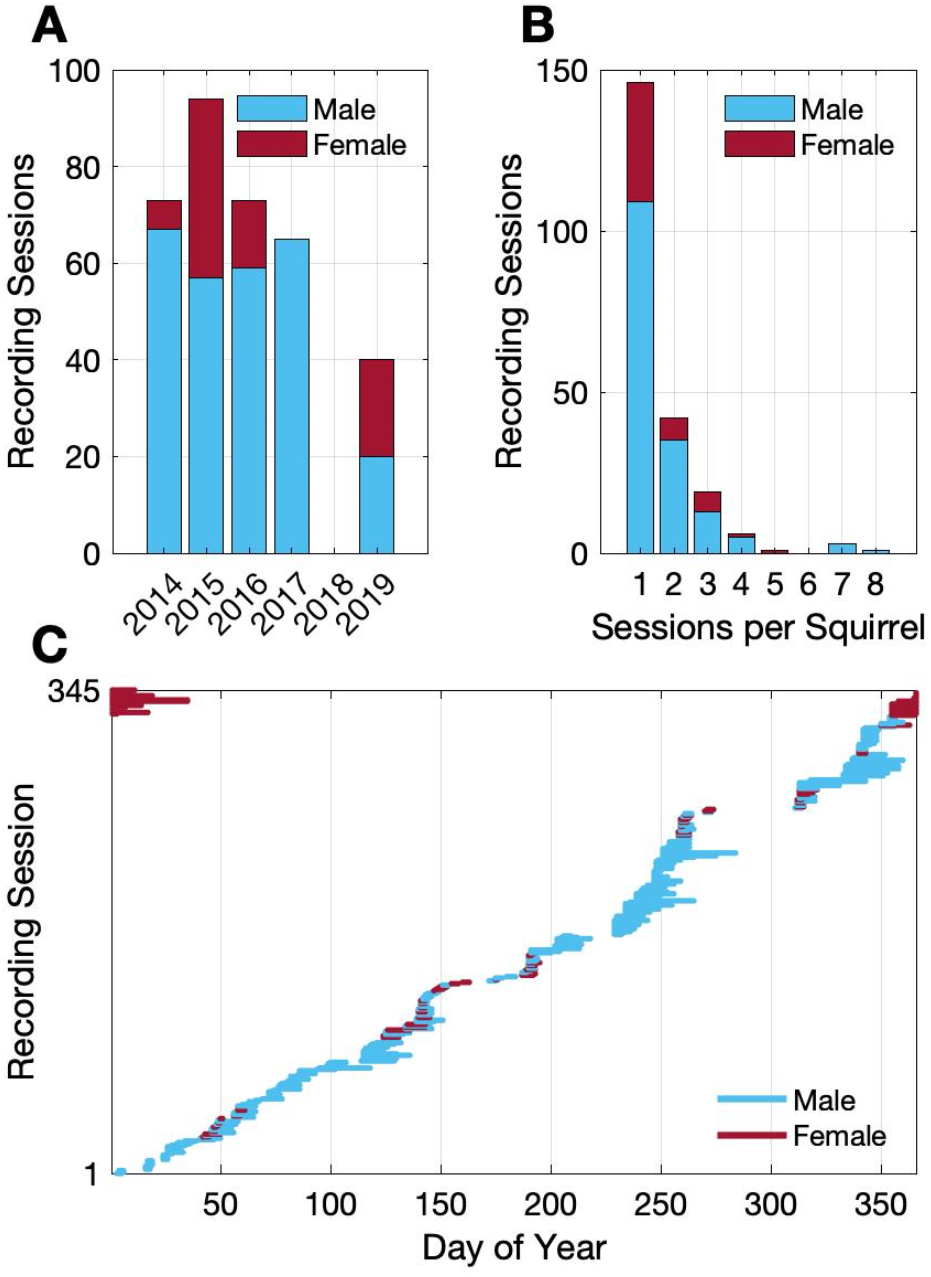
Data Overview. (A) Recording sessions by year (268 male, 77 female). Lactating and pregnant females are not present in these data. (B) Number of recording sessions that are from the same squirrel (n = 218 total squirrels). For example, the same squirrel may have been recorded in 2016 and 2017, contributing to the magnitude of the bar above x-axis ‘2’. (C) Recording duration for each recording session (n = 345 total), sorted by start day.

**Figure S2.**
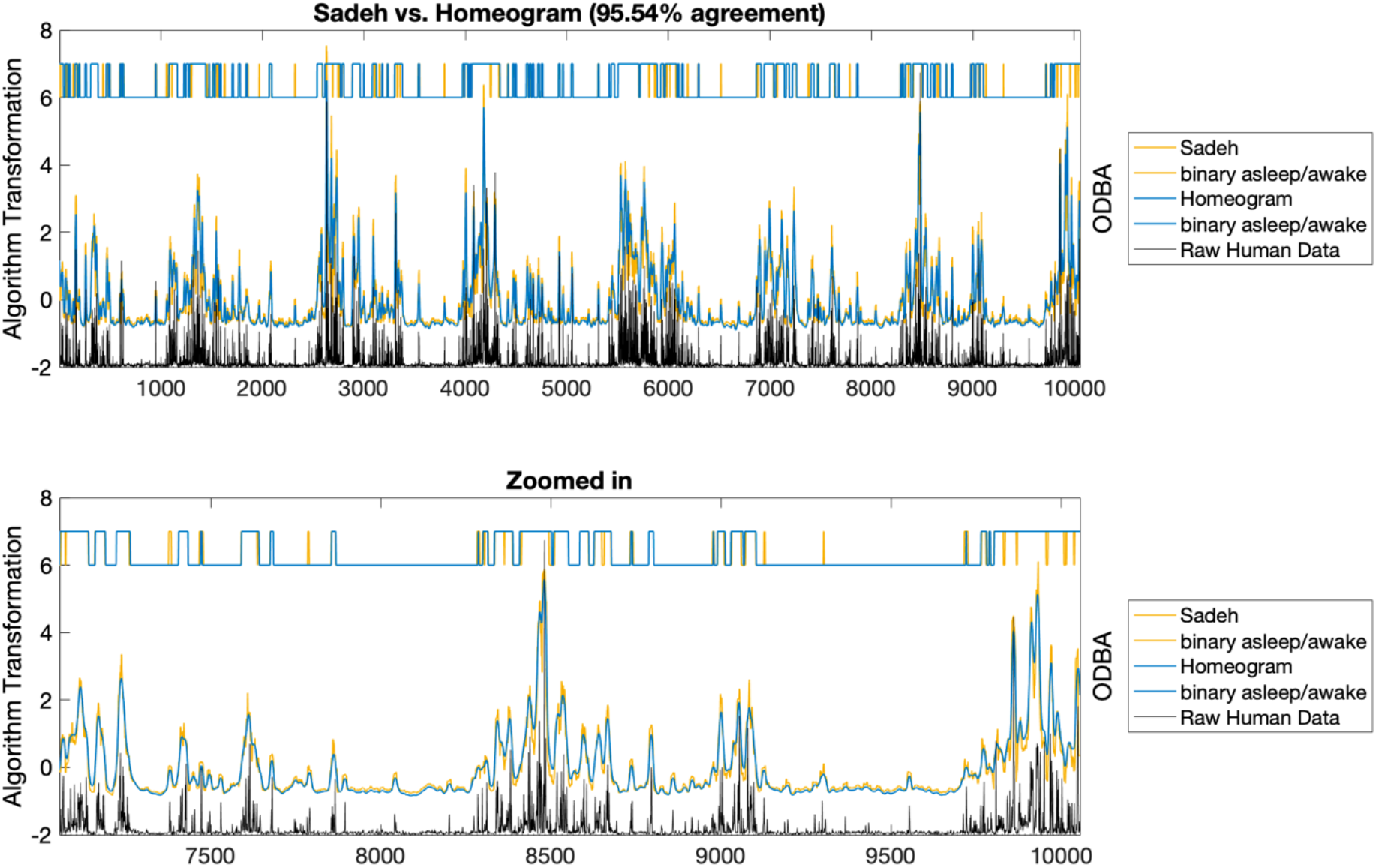
Homeogram agreement with conventional algorithm. (Top) Sadeh algorithm (yellow) and Homeogram (blue) methods applied to 7 days of human actigraphy (black, right ODBA axis). The algorithmic transformation is shown in analog form (left y-axis) and then binarized when the probability for asleep state is greater than 0, offset from the left-axis between values 6-7. (Bottom) Zoomed in version.

**Table S1.** Sleep statistics for North American red squirrels in Yukon, Canada derived from homeogram using accelerometry data from only non-mast years (2015, 2016, 2017) and mast years (2014, 2019).

Non-Mast Year
|  | Winter | Spring | Summer | Autumn | All |
| --- | --- | --- | --- | --- | --- |
| Day Length (hrs.) | $6.57 \pm 0.82$ | $12.07 \pm 2.52$ | $18.20 \pm 0.82$ | $12.74 \pm 2.46$ | 12 |
| Sleep per day (hrs.) | $10.49 \pm 1.90$ | $9.32 \pm 2.41$ | $8.30 \pm 1.88$ | $8.95 \pm 1.61$ | $9.66 \pm 2.15$ |
| Sleep in daylight (hrs.) | $0.72 \pm 0.86$ | $1.57 \pm 1.73$ | $3.21 \pm 1.87$ | $1.13 \pm 1.29$ | $1.34 \pm 1.59$ |
| Sleep in darkness (hrs.) | $9.76 \pm 1.82$ | $7.75 \pm 1.73$ | $5.08 \pm 0.98$ | $7.82 \pm 1.09$ | $8.31 \pm 2.28$ |
| Total sleep transitions | $28 \pm 7$ | $22 \pm 8$ | $17 \pm 7$ | $16 \pm 6$ | $23 \pm 9$ |
| Sleep transitions in daylight | $3 \pm 3$ | $6 \pm 4$ | $11 \pm 5$ | $5 \pm 4$ | $5 \pm 5$ |
| Sleep transitions in darkness | $26 \pm 6$ | $17 \pm 8$ | $6 \pm 4$ | $11 \pm 5$ | $18 \pm 9$ |

Mast Year
|  | Winter | Spring | Summer | Autumn | All |
| --- | --- | --- | --- | --- | --- |
| Day Length (hrs.) | $6.57 \pm 0.82$ | $12.07 \pm 2.52$ | $18.20 \pm 0.82$ | $12.74 \pm 2.46$ | 12 |
| Sleep per day (hrs.) | NaN ± NaN | $10.19 \pm 1.60$ | $8.83 \pm 2.05$ | $8.50 \pm 1.75$ | $9.05 \pm 1.98$ |
| Sleep in daylight (hrs.) | NaN ± NaN | $2.29 \pm 1.50$ | $4.47 \pm 2.01$ | $0.71 \pm 1.68$ | $2.95 \pm 2.44$ |
| Sleep in darkness (hrs.) | NaN ± NaN | $7.90 \pm 1.82$ | $4.36 \pm 1.06$ | $7.79 \pm 1.38$ | $6.10 \pm 2.21$ |
| Total sleep transitions | NaN ± NaN | $26 \pm 7$ | $20 \pm 7$ | $13 \pm 6$ | $19 \pm 8$ |
| Sleep transitions in daylight | NaN ± NaN | $7 \pm 4$ | $13 \pm 6$ | $3 \pm 5$ | $9 \pm 7$ |
| Sleep transitions in darkness | NaN ± NaN | $19 \pm 7$ | $7 \pm 4$ | $10 \pm 4$ | $10 \pm 7$ |

**Table S2.**
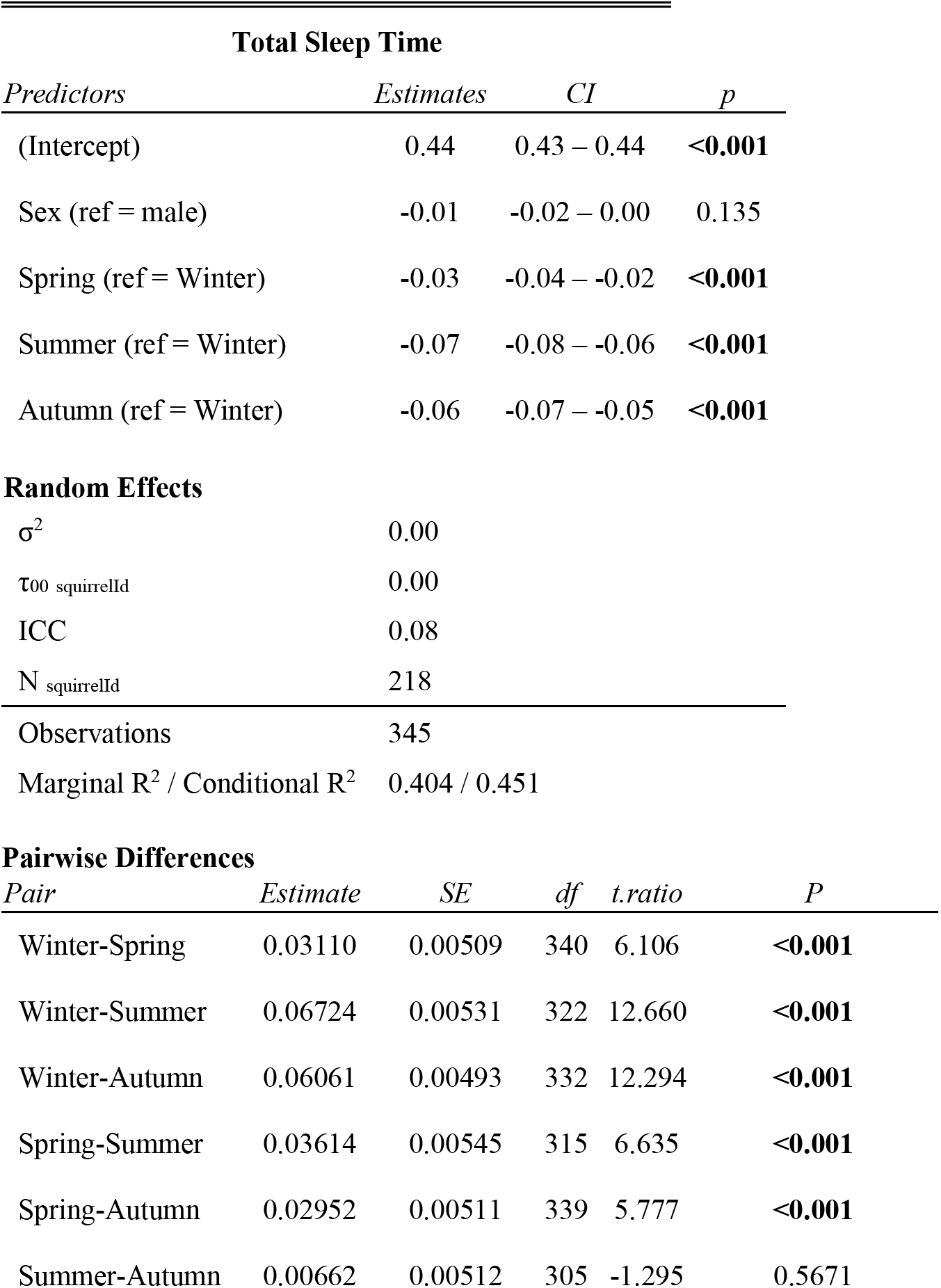
Are there differences in total sleep time? Results from linear mixed-effects models followed by pairwise comparisons to examine if total sleep time varied according to sex or season.

**Table S3.**
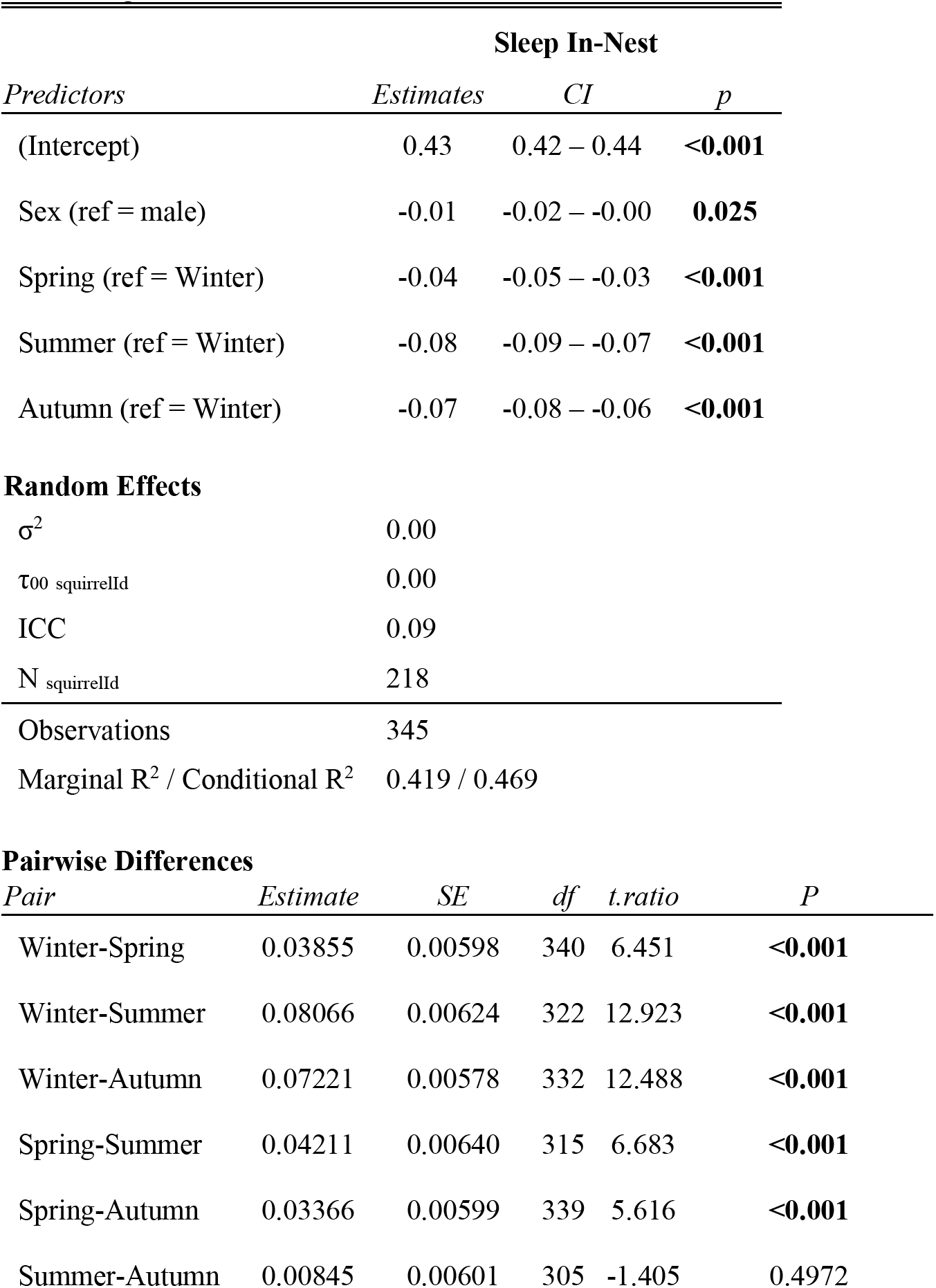
Are there differences in the time slept inside the nest? Results from linear mixed-effects models followed by pairwise comparisons to examine if sleep time inside the nest varied according to sex or season.

**Table S4.** Are there differences in the time slept outside the nest? Results from linear mixed-effects models followed by pairwise comparisons to examine if sleep time outside the nest varied according to sex or season.

| Sleep Out of Nest |  |  |  |  |  |
| --- | --- | --- | --- | --- | --- |
| Predictors | Estimates | CI | p |  |  |
| (Intercept) | 0.01 | 0.00 – 0.02 | **0.001** |  |  |
| Sex (ref = male) | 0.01 | -0.00 – 0.01 | 0.149 |  |  |
| Spring (ref = Winter) | 0.01 | -0.00 – 0.01 | 0.081 |  |  |
| Summer (ref = Winter) | 0.01 | 0.01 – 0.02 | **0.001** |  |  |
| Autumn (ref = Winter) | 0.01 | 0.00 – 0.02 | **0.003** |  |  |
| **Random Effects** |  |  |  |  |  |
| $\sigma^2$ | 0.00 | | | | |
| $\tau_{00\_squirrelId}$ | 0.00 | | | | |
| ICC | 0.05 |  |  |  |  |
| N $\_squirrelId$ | 218 | | | | |
| Observations | 345 |  |  |  |  |
| Marginal R<sup>2</sup> / Conditional R<sup>2</sup> | 0.049 / 0.095 |  |  |  |  |
| **Pairwise Differences** |  |  |  |  |  |
| Pair | Estimate | SE | df | t.ratio | P |
| Winter-Spring | -0.00699 | 0.00401 | 340 | -1.743 | 0.3031 |
| Winter-Summer | -0.01338 | 0.00417 | 319 | -3.212 | **0.0079** |
| Winter-Autumn | -0.01136 | 0.00389 | 334 | -2.921 | **0.0194** |
| Spring-Summer | -0.00639 | 0.00427 | 309 | -1.498 | 0.4402 |
| Spring-Autumn | -0.00437 | 0.00402 | 340 | -1.087 | 0.6976 |
| Summer-Autumn | 0.00202 | 0.00401 | 297 | 0.504 | 0.9581 |

**Table S5.** Are there differences in the proportion of time slept in the nest? Results from linear mixed-effects model to examine if proportion of sleep (sleep in-nest / total sleep) varied according to sex or season.

| Proportion of Sleep In-Nest |  |  |  |  |  |
| --- | --- | --- | --- | --- | --- |
| Predictors | Estimates | CI | p |  |  |
| (Intercept) | 0.52 | 0.51 – 0.54 | <0.001 |  |  |
| Sex (ref = male) | -0.01 | -0.03 – 0.01 | 0.206 |  |  |
| Spring (ref = Winter) | 0.06 | 0.04 – 0.08 | <0.001 |  |  |
| Summer (ref = Winter) | 0.07 | 0.05 – 0.09 | <0.001 |  |  |
| Autumn (ref = Winter) | 0.14 | 0.12 – 0.16 | <0.001 |  |  |
| Random Effects |  |  |  |  |  |
| $\sigma^2$ | 0.00 | | | | |
| $\tau_{00\_squirrelId}$ | 0.00 | | | | |
| ICC | 0.15 |  |  |  |  |
| N $\_squirrelId$ | 218 | | | | |
| Observations | 345 |  |  |  |  |
| Marginal R<sup>2</sup> / Conditional R<sup>2</sup> | 0.349 / 0.449 |  |  |  |  |
| Pairwise Differences |  |  |  |  |  |
| Pair | Estimate | SE | df | t.ratio | P |
| Winter-Spring | -0.0593 | 0.0104 | 338 | -5.710 | <0.001 |
| Winter-Summer | -0.0702 | 0.0109 | 327 | -6.419 | <0.001 |
| Winter-Autumn | -0.1375 | 0.0100 | 327 | -13.745 | <0.001 |
| Spring-Summer | -0.0109 | 0.0112 | 324 | -0.971 | 0.7663 |
| Spring-Autumn | -0.0782 | 0.0104 | 335 | -7.516 | <0.001 |
| Summer-Autumn | -0.0673 | 0.0106 | 317 | -6.366 | <0.001 |

**Table S6.**
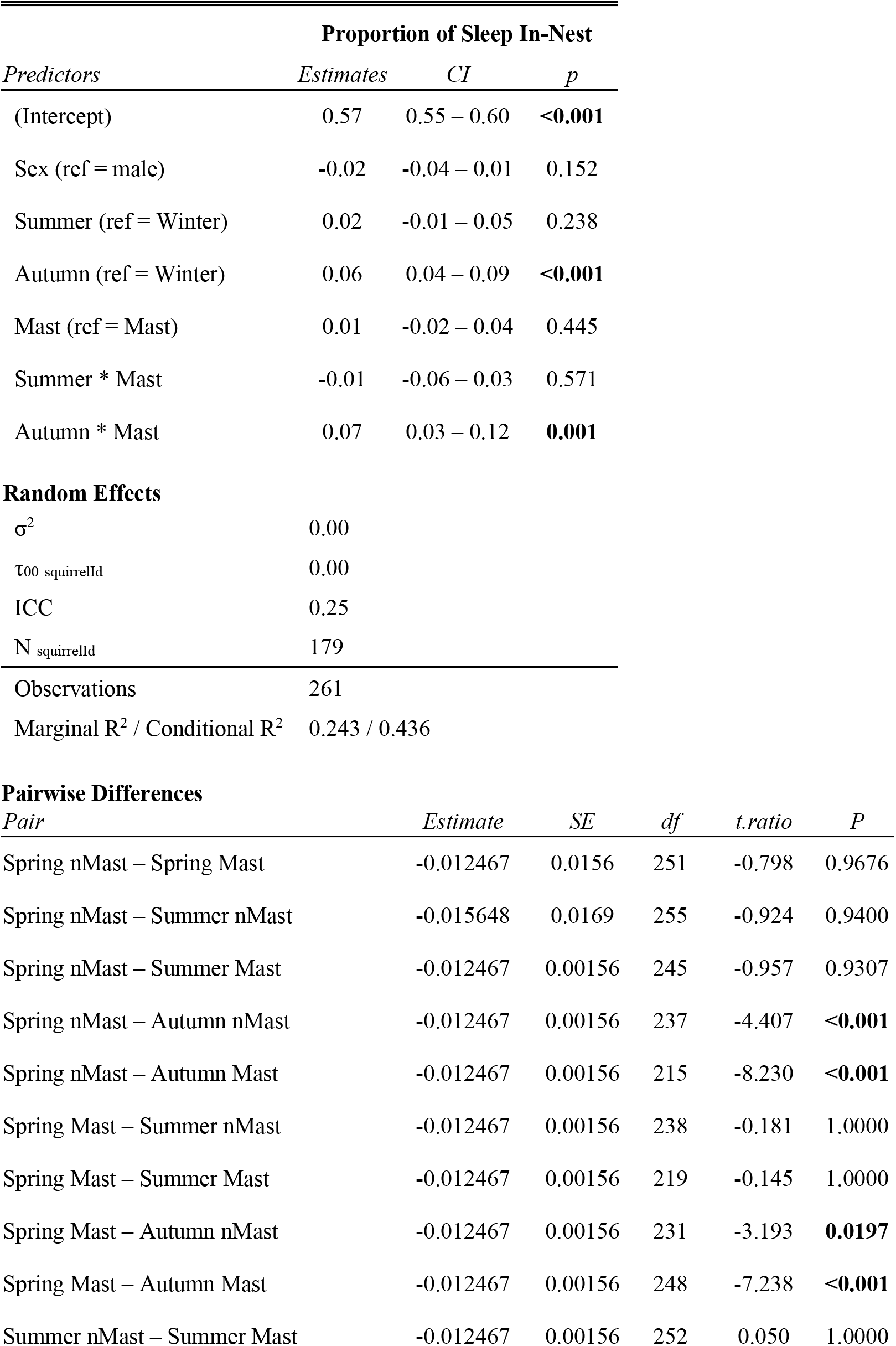
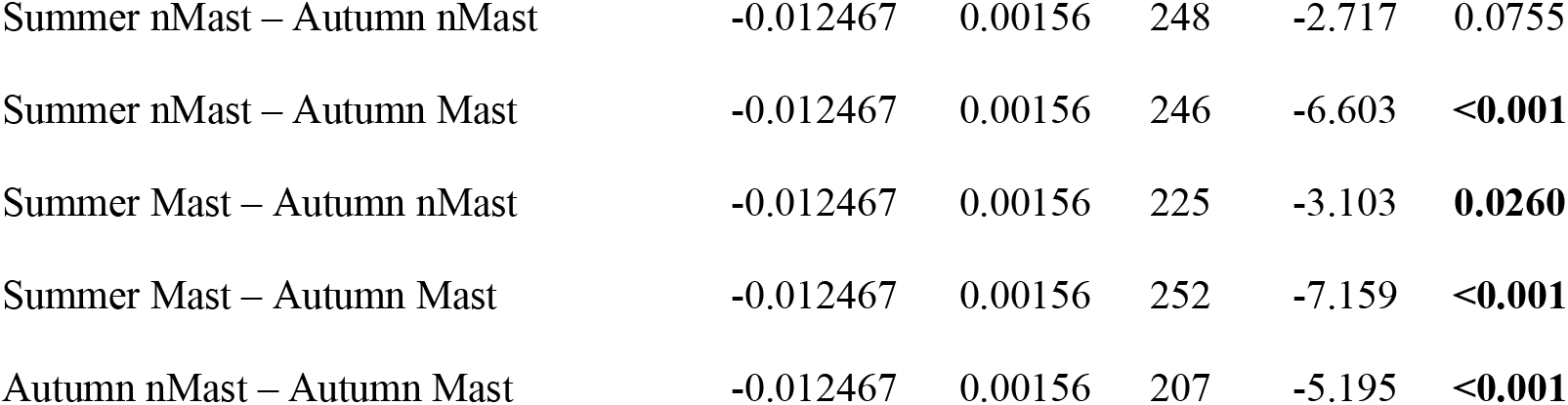
Are there interaction effects between season and mast that change the proportion of time slept in the nest? Results from linear mixed-effects models followed by pairwise comparisons to examine if proportion of sleep (sleep in-nest / total sleep) varied according to sex or season.

| *Predictors* | **Proportion of Sleep In-Nest** |  |  |
| --- | --- | --- | --- |
|  | *Estimates* | *CI* | *p* |
| (Intercept) | 0.57 | 0.55 – 0.60 | **<0.001** |
| Sex (ref = male) | -0.02 | -0.04 – 0.01 | 0.152 |
| Summer (ref = Winter) | 0.02 | -0.01 – 0.05 | 0.238 |
| Autumn (ref = Winter) | 0.06 | 0.04 – 0.09 | **<0.001** |
| Mast (ref = Mast) | 0.01 | -0.02 – 0.04 | 0.445 |
| Summer * Mast | -0.01 | -0.06 – 0.03 | 0.571 |
| Autumn * Mast | 0.07 | 0.03 – 0.12 | **0.001** |

Random Effects
|  |  |
| --- | --- |
| $\sigma^2$ | 0.00 |
| $\tau_{00\_squirrelId}$ | 0.00 |
| ICC | 0.25 |
| $N_{squirrelId}$ | 179 |
| Observations | 261 |
| Marginal $R^2$ / Conditional $R^2$ | 0.243 / 0.436 |

Pairwise Differences
| *Pair* | *Estimate* | *SE* | *df* | *t.ratio* | *P* |
| --- | --- | --- | --- | --- | --- |
| Spring nMast – Spring Mast | -0.012467 | 0.0156 | 251 | -0.798 | 0.9676 |
| Spring nMast – Summer nMast | -0.015648 | 0.0169 | 255 | -0.924 | 0.9400 |
| Spring nMast – Summer Mast | -0.012467 | 0.00156 | 245 | -0.957 | 0.9307 |
| Spring nMast – Autumn nMast | -0.012467 | 0.00156 | 237 | -4.407 | **<0.001** |
| Spring nMast – Autumn Mast | -0.012467 | 0.00156 | 215 | -8.230 | **<0.001** |
| Spring Mast – Summer nMast | -0.012467 | 0.00156 | 238 | -0.181 | 1.0000 |
| Spring Mast – Summer Mast | -0.012467 | 0.00156 | 219 | -0.145 | 1.0000 |
| Spring Mast – Autumn nMast | -0.012467 | 0.00156 | 231 | -3.193 | **0.0197** |
| Spring Mast – Autumn Mast | -0.012467 | 0.00156 | 248 | -7.238 | **<0.001** |
| Summer nMast – Summer Mast | -0.012467 | 0.00156 | 252 | 0.050 | 1.0000 |
| Summer nMast – Autumn nMast | -0.012467 | 0.00156 | 248 | -2.717 | 0.0755 |
| Summer nMast – Autumn Mast | -0.012467 | 0.00156 | 246 | -6.603 | **<0.001** |
| Summer Mast – Autumn nMast | -0.012467 | 0.00156 | 225 | -3.103 | **0.0260** |
| Summer Mast – Autumn Mast | -0.012467 | 0.00156 | 252 | -7.159 | **<0.001** |
| Autumn nMast – Autumn Mast | -0.012467 | 0.00156 | 207 | -5.195 | **<0.001** |

